# Huntingtin S421 phosphorylation increases kinesin and dynein engagement on early endosomes and lysosomes

**DOI:** 10.1101/2022.05.27.493751

**Authors:** Emily N. P. Prowse, Abdullah R. Chaudhary, David Sharon, Adam G. Hendricks

## Abstract

Huntingtin (HTT) is a scaffolding protein that recruits motor proteins to vesicular cargoes, enabling it to regulate kinesin-1, dynein, and myosin-VI-dependent transport. To maintain the native stoichiometry of huntingtin with its interacting partners, we used CRISPR/Cas9 to induce a phosphomimetic mutation of the endogenous HTT at S421 (HTT-S421D). Using single particle tracking, optical tweezers, and immunofluorescence, we examined the effects of this mutation on the motility of early endosomes and lysosomes. In HTT-S421D cells, lysosomes exhibit longer displacements and higher processive fractions compared to wild-type (HTT-WT) cells. Kinesins and dyneins exert greater forces on early endosomes and lysosomes in cells expressing HTT-S421D. Additionally, endosomes bind to microtubules faster and are more resistant to detachment under load. The recruitment of kinesins and dyneins to microtubules is enhanced in HTT-S421D cells. In contrast, overexpression of HTT had variable effects on the processivity, displacement, and directional bias of both early endosomes and lysosomes. These data indicate that phosphorylation of the endogenous huntingtin causes early endosomes and lysosomes to move longer distances and more processively by recruiting and activating both kinesin-1 and dynein.

**Statement of Significance:** The ubiquitous scaffolding protein huntingtin regulates the recruitment and activity of microtubule motors. Huntingtin phosphorylation at S421 enhances the microtubule binding and force generation of kinesin and dynein on early endosomes and lysosomes. Using optical tweezers to measure the forces exerted on endosomes in CRISPR-engineered cells, we find that a phosphomimetic huntingtin mutation (S421D) enhances both kinesin- and dynein-driven forces on early endosomes and lysosomes. The ability to modulate motor activity on a range of organelles makes huntingtin unique and suggests a significant role for huntingtin in regulating intracellular transport.

## INTRODUCTION

Huntingtin (HTT) is an essential, multifunctional, and ubiquitous protein. It consists of several disordered regions and 34 α-helical HEAT repeats (**Figure 1a**) (1). HTT is essential for embryonic development (2–5), and is ubiquitously expressed in tissues including the brain, ovaries, and testis into adulthood (6, 7). It has diverse roles within the cell, including the regulation of gene expression (8–13), cell morphology (14, 15), cell survival (16), and intracellular transport (17–22). HTT enhances transport of brain derived neurotrophic factor (BDNF) vesicles (17, 22, 23), autophagosomes in the mid-axon (18, 19), TrkB signalling vesicles (20), amyloid precursor protein (APP) vesicles (24), and synaptic vesicles through GABA_R_ signalling (21). The remarkable ubiquity and variety of HTT’s roles underscore the importance of understanding its function in health and disease.

**Figure 1.**
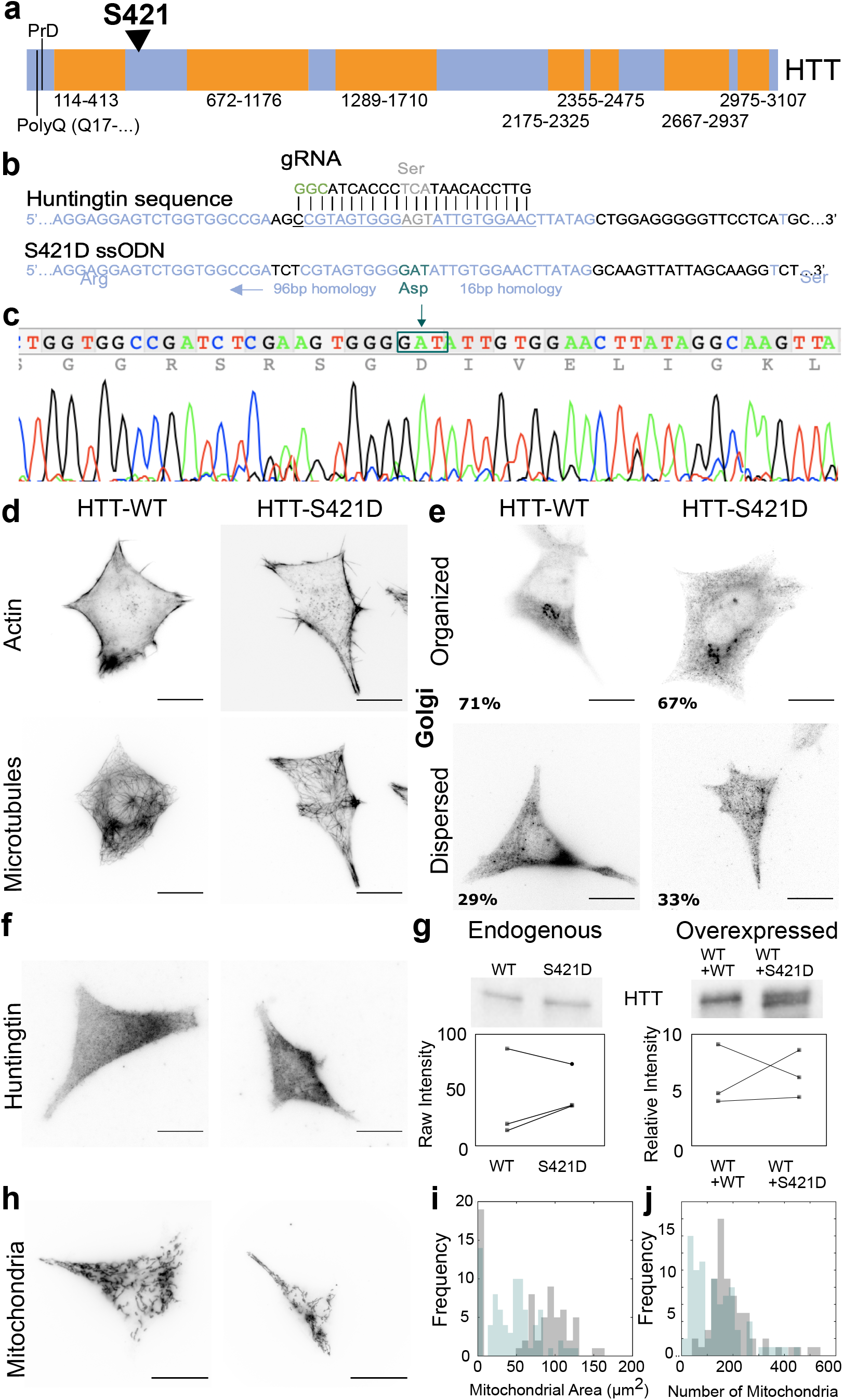
HTT-S421D HEK293T cell line does not affect cytoskeletal organization, huntingtin localization, or dynein function. (**a**) Huntingtin protein sequence schematic demonstrating the structured domains (HEAT repeats and other α-helical regions, orange), unstructured domains (blue), S421 phosphosite (black arrowhead), polyglutamine repeat region (PolyQ), and proline rich domain (PrD). (**b**) CRISPR phosphomimetic mutant design guide RNA (gRNA), and ssODN homologous recombination template. (**c**) Chromatogram of the positive clone containing the phosphomimetic mutation HTT-S421D. (**d**) Immunostaining of actin (top) and microtubules (bottom) indicating no difference between HTT-WT (left) and HTT-S421D (right) HEK293T cells. Note immunofluorescence images are inverted such that the protein of interest is dark, and the background is white, and all scale bars in the figure are 20 μm. (**e**) In both HTT-WT and HTT-S421D HEK293T cells, the Golgi apparatus appeared compact (organized) in ~70% of cells and dispersed in ~30% of cells. HTT-WT data for actin, microtubules, and Golgi apparatus staining was from 14 experiments, 84 cells, and HTT-S421D data from 11 experiments, 69 cells. (**f**) Immunofluorescence of huntingtin protein localization in HTT-WT (left) compared to HTT-S421D (right) HEK293T cells. Immunofluorescence of huntingtin localization for HTT-WT cells from 3 experiments, 43 cells and for HTT-S421D from 2 experiments, 22 cells. (**g**) HTT-WT and HTT-S421D huntingtin endogenous expression levels (top left) detected by Western blot with the raw intensity data from three independent samples (bottom left), compared to levels when overexpressing full length mCherry-HTT-WT (WT+WT) or mCherry-HTT-S421D (WT+S421D) huntingtin (top right) and the fold change relative to native expression levels for three independent samples (bottom right). Samples collected on the same date are connected by a line. (**h**) Mitochondrial morphology of HTT-WT (left) and HTT-S421D (right) HEK293T cells. (**i**) Mitochondrial area of HTT-WT (grey) compared to HTT-S421D (blue) cells, p=6.59×10^−8^ by Kolmogorov-Smirnov (ks) test. (**j**) Number of mitochondria per cell in HTT-WT (grey) compared to HTT-S421D (blue), p=6.14×10^−7^ by ks test. Mitochondrial analysis for HTT-WT from 7 experiments, 88 cells, and HTT-S421D from 7 experiments, 94 cells.

Teams of kinesins and dyneins transport signalling and degradative cargoes. There are multiple kinesins that drive outward transport towards microtubule plus ends at the cell periphery. Different ratios of kinesin-1, kinesin-2, and kinesin-3 associate with distinct cargo populations, allowing different motility characteristics for each population of vesicles (25). Each type of kinesin has a unique binding rate and velocity, which leads combinations of motors to create a motility barcode for cargoes. Unlike kinesins, there is only one form of cytoplasmic dynein that drives organelle transport. Dynein forms a complex with dynactin and activating adaptors such as BicD, Hook1, and others to direct inward transport towards microtubule minus ends at the cell center (19, 26, 27). Scaffolding and adaptor proteins work together to regulate the recruitment and activity of both kinesin and dynein motors on each cargo based on its identity.

Huntingtin recruits teams of kinesin, dynein, and myosin VI motors to cargoes, allowing it to control the cargo’s direction of movement along microtubules and switching between actin and microtubules (28). HTT interacts with molecular motors indirectly through adaptors; kinesin-1 and the dynein adaptor dynactin interact with HTT through HAP1 (21, 29–31), and myosin VI interacts through optineurin (32). In addition, dynein interacts directly with HTT through its intermediate chain, suggesting a prominent role for HTT in regulating inward transport (33). HTT’s interaction with myosin VI suggests HTT regulates transitions to the actin cytoskeleton for short range transport or tethering. Upon HTT deletion or mutation, intracellular transport of many vesicular cargoes is severely inhibited (34, 35).

Polyglutamine expansion mutations in HTT lead to Huntington’s disease (HD), disrupting multiple cellular functions and leading to neurodegeneration in medium spiny neurons of the striatum (36). HD mutations result in aberrant initiation of autophagy, mTOR signalling, mitochondrial fission, and palmitoylation (reviewed in (37)). Additional defects occur in neuronal transcription, endoplasmic reticulum homeostasis, mitochondrial function, calcium signalling, synaptic activity, and intracellular transport (reviewed in (38)) demonstrating the importance of HTT’s native interaction network in maintaining homeostasis (23, 39, 40). HD mutations disrupt tight junctions and endocytosis leading to increased metastasis in breast cancer (15, 41); further, huntingtin phosphorylation at S421 is decreased in cancer cells and blocking HTT phosphorylation results in enhanced metastasis (15). Thus, huntingtin dysfunction plays roles in both neurodegeneration and cancer.

Huntingtin is regulated by multiple post-translational modifications, the functions, and mechanisms of which remain unknown. Its palmitoylation site at C214, regulated by HIP14 in the Golgi, facilitates interactions with vesicular membranes (42). Palmitoylation is reduced in cells expressing the polyglutamine-expanded HTT protein that leads to HD, while restoring palmitoylation reduces aggregation and cytotoxicity in vitro (40). Sumoylation sites may contribute to nuclear import and transcriptional regulation (43). Ubiquitination and acetylation regulate huntingtin-mediated autophagy (18).

HTT has 40 known phosphosites (reviewed in (38)). In particular, the phosphorylation state of S421 affects axonal transport direction and velocity of BDNF vesicles and autophagosomes (18, 39), while mitochondrial transport is unaffected by this phosphosite (44). The phosphomimetic mutation (HTT-S421D) mimics a constitutively phosphorylated amino acid and biased BDNF velocity and run length transport towards the cell periphery, while phosphoresistive mutations (HTT-S421A) led to increased transport towards the cell center (22, 39). In primary neurons derived from mice with the HTT-S421D and HTT-S421A mutation, BDNF vesicles and APP vesicles demonstrated an outward directional bias with increased recruitment of kinesin to APP vesicles upon S421 phosphorylation (22, 24). The HTT-S421D mutation is neuroprotective in cells with the mutation that leads to HD (23), which motivates our attempt to understand the native role of this phosphosite in cells with normal (WT) HTT. It has also been generalized for various scaffolding proteins including RILP, Miro/TRAK1/TRAK2, JIP1, JIP3/4 that post-translational modifications can modify transport direction (45). These studies suggest that huntingtin phosphorylation directs transport towards the cell periphery.

Previous studies analyzed the motility of cargoes overexpressing HTT-S421D or HTT-S421A constructs in COS7, HEK, and primary rat neurons and performed Western blotting to determine S421’s role in transport regulation (23, 39). Western blot and immunofluorescence in overexpressing COS7 cells suggest that S421 phosphorylation increases the recruitment of kinesin-1 to cargoes (23, 39). However, only a subset of the motors bound to a cargo might be active (46). In addition, the stoichiometry of interactions between scaffolding proteins, adaptors, and motors is disrupted in overexpression systems. We mutated the endogenous HTT gene to retain HTT’s native interactions and expression levels and observed its effect on organelle transport. Additionally, we measured the forces generated by the motors driving early endosome and lysosome transport with live cell optical tweezers assays to determine whether phosphorylation of huntingtin modifies motor activity. Combining biophysical approaches with gene editing provided mechanistic insight into HTT’s regulation of intracellular transport by S421 phosphorylation.

To understand how S421 phosphorylation of HTT regulates intracellular transport, we examined early endosome and lysosome transport in CRISPR-generated HTT-S421D HEK293T cells. Motility assays demonstrate no change in directional bias for early endosomes nor lysosomes in cells expressing HTT-S421D. We show that early endosomes’ processive fraction slightly increased, while lysosome displacement and processive fraction increased upon S421 pseudo-phosphorylation. Examining the forces generated by microtubule motors on early endosomes and lysosomes, we observed both kinesin and dynein motors had higher binding rates and increased resistance to unbinding under load with HTT-S421D for both cargoes. In cells expressing HTT-S421D, we show that kinesins and dyneins exert higher forces in both outward and inward directions on early endosomes and lysosomes. Adaptors such as JIP1 (48), TRAK2 (47, 49), and Hook3 (50) act as activating adaptors for both kinesins and dyneins, however not simultaneously. Here, we show that huntingtin acts as a scaffolding protein for both kinesin and dynein and enhances force generation of both motors simultaneously in living cells. Activation of kinesin and dynein led to enhanced displacement and processive fraction of endocytic cargoes. Further, we show higher kinesin and dynein forces have differential effects on the motility of early endosomes and lysosomes. By immunofluorescence, a higher fraction of kinesins and dyneins are associated with microtubules upon S421 phosphorylation. The phosphorylation-induced change in force generation for lysosomes was more subtle than for early endosomes. We hypothesize that huntingtin regulates transport similarly for early endosomes and lysosomes, however its effects depend on the cargo’s identity and motility characteristics. HTT phosphorylation recruits and activates kinesin-1 and dynein teams, leading to a pronounced increase in processive fraction and displacement for lysosomes, while the effect on the motility of early endosomes is more subtle. Our data indicates phosphorylation of HTT at S421 increases motor recruitment on and activity of early endosomes and lysosomes. Additionally, overexpression of WT+HTT-WT and WT+HTT-S421D suggests overexpressed HTT disrupts the transport complex of HTT, motors, and adaptors required for transport, affecting cargo displacement, processivity, and direction.

## MATERIALS AND METHODS

### Cell Culture

We passaged HEK293T cells (donated by Dr. Amine Kamen’s lab) at 80% or higher confluency in a biosafety cabinet starting by rinsing with 2.5 mL 37 °C 1X phosphate buffered saline (PBS, Wisent, St-Jean-Baptiste, Quebec), followed by adding 400 μL of 4 °C 0.25% trypsin (Wisent) and incubating for 5 mins at 37 °C. Next, we used 5 mL of 37 °C complete medium (Dulbecco’s Modified Eagle Medium (DMEM), 10% fetal bovine serum (FBS), 2 mM Glutamax, 60 μg/mL penicillin 100 μg/mL streptomycin, all from ThermoFisher Scientific, Waltham, Massachusetts) to neutralize the trypsin activity. Finally, we placed a 1/5-1/20 dilution of the resuspended cells in 37 °C fresh complete medium and transferred to the 37 °C, 5% CO2 incubator. We tested all cell stocks for mycoplasma.

### CRISPR Design and Molecular cloning

We transfected HEK293T cells with 1.25 μg of lentiCRISPR v2 cassette and 1.25 μg of the single-stranded oligodeoxynucleotide (ssODN) for 24-36 hrs. lentiCRISPR v2 was a gift from Feng Zhang (Addgene plasmid # 52961; http://n2t.net/addgene:52961; RRID:Addgene_52961) (51). Subsequently, we selected expressing cells using 2 μg puromycin for 48-72 hrs. Finally, we isolated single cells using fluorescence activated single cell sorting and grew them for 2-3 weeks in complete media containing 50% FBS. We verified clones containing the S421D sequence with next-generation sequencing.

gRNA sequence:
GTTCCACAATACTCCCACTACGG
ssODN sequence:
ACGCCTCCACCCGAGCTTCTGCAAACCCTGACCGCAGTCGGGGGCATTGGGCAGCTCACCGCTGCTAAGGAGGAGTCTGGTGGCCGATCTCGTAGTGGGGATATTGTGGAACTTATAGGCAAGTTATTAGCAAGGTCTACTCTTACAATTAACTTTGCAGTAATACTAGTTACACTCTATTGATTATGGGCCTGCCCTGT
Sequencing primers:
Forward: CTCATGCTGCTCCTCTAGTGCA
Reverse: GGGGCTTTAAATGTGTGGCCTC

### Cell Ploidy Analysis

We trypsinized both cell lines as described in the cell culture methods, centrifuged them for 5 mins at 1200 rpm in 5 mL of DMEM complete, and resuspended them in 37°C 4% paraformaldehyde (Millipore Sigma, Burlington, Massachusetts) in PBS (Wisent) for 15 minutes. We then centrifuged the cells for 5 mins at 1200 rpm and resuspended them in PBS, such that there were approximately 12 000 000 cells/mL with a minimum volume of 300 μL. The cells were stored at 4°C in PBS. We added 14.3 nM DAPI (D1306, ThermoFisher Scientific) to the cells 20 mins prior to FACS. The FACS technician then performed the cell sorting, isolating the data for single cells and determining the DAPI intensity from each cell.

### Plasmids and Transfection

We transfected cells with 0.8 μg of eGFP-Rab5 or mCherry-S421D-HTT or mCherry-WT-HTT constructs 24-48 hrs before imaging. mCherry-S421D-HTT and mCherry-WT-HTT constructs were generously donated from the Saudou lab, generated from the full length pARIS-HTT (52) using methods from their previous work (53). We replaced cell culture medium in the 35 mm round glass bottom dishes containing the cells with 0.2 mL serum-free DMEM. We separately diluted 2 μL/dish Lipofectamine 2000 (ThermoFisher Scientific) to 50 μL/dish serum-free DMEM and 0.8 μg/dish of DNA to 50 μL/dish of serum-free DMEM. We then mixed the DNA and lipofectamine, incubated for 20 mins at room temperature, and added 100 μL/dish to the cells. Four hours later, we replaced the serum-free medium with complete media.

### Western Blotting

We lysed cells using a Dounce homogenizer 20-30 times in 400 μL PBS (Wisent) containing 0.01 M dithiothreitol (DTT, ThermoFisher Scientific) and 1X protease inhibitor cocktail (100X stock, BioShop, Burlington, Ontario) on ice. Subsequently, we centrifuged the cell solution at 1500 g for 10 mins at 4 °C, then centrifuged the supernatant at 16000 g for 10 mins at 4 °C. We heated samples to diluted in diluted in 2x Laemmli (BioRad, Hercules, California) with 2.5% 2-mercaptoethanol (Millipore Sigma) 100 °C for at least 10 mins before loading added 20 μL of lysate to the SDS-PAGE gel. We used a 25 mM Tris (BioShop), 190 mM glycine (Wisent), and 0.1% SDS (BioShop) running buffer, and applied 80 mA current for 60-90 mins. To transfer, we used 25 mM Tris (BioShop), 190 mM glycine (BioShop), and 30% methanol (Millipore Sigma) transfer buffer for wet transfer at 100 V for 90 mins with a PVDF membrane. We blocked the membrane for 1 hour with blocking buffer: 5% bovine serum albumin (BSA, BioShop) in tris-buffered saline (BioShop) with Tween 20 (Millipore Sigma) (TBST, 20 mM Tris, 150 mM NaCl, 0.1% Tween 20). We diluted primary antibodies in the blocking buffer and incubated at 4 °C overnight. Next, we washed the membrane with blocking buffer 3 times for 5 mins and incubated with the secondary antibody diluted in blocking buffer for 90 mins with constant agitation. We washed the membrane 3 times with TBST before incubating with a 1:1 ratio of horseradish peroxidase substrate and enzyme mixture (Millipore, Burlington, Massachusetts). We imaged using a Biorad Chemidoc camera set to detect chemiluminescence. We subsequently stained the membrane with Coomassie G-250 (BioShop), using a previously established protocol from Goldman et al. titled Coomassie blue R-250 staining (54). We used the following primary antibodies: 1/1000 dilution of T9026 ms α-tubulin, (Millipore Sigma lot: 000089497) and 1/500 dilution of MAB2166 ms-huntingtin, (Abcam, Cambridge, UK lot: 3703568). The secondary antibody was rabbit anti-mouse ab6728 (Abcam).

### Motility Assay

In a biological safety cabinet, we mixed 200 μL of 37 °C complete medium with LysoTracker™ Deep Red (ThermoFisher Scientific) to a concentration of 70 nM and added it to HEK293T cells in MatTek 35 mm glass bottom dishes. Cells were incubated for 10 mins at 37 °C, and subsequently we replaced the medium with 1 mL of 37 °C Lebovitz (ThermoFisher Scientific) with 10 % FBS for imaging. We imaged cells in a 37°C chamber of a Nikon Eclipse Ti-E microscope in near-total internal reflection (TIRF) fluorescence settings using an EMCCD camera with an exposure time of 120 ms for 3 mins. We repeated the process for several cells within each dish and all cells were imaged within 1 hr of adding Lysotracker. For Rab5 transfected cells, we replaced the medium with 0.5-1 mL of 37 °C Lebovitz with 10 % FBS and imaged them as described for the lysosomes. For experiments using 250 nM latrunculin A (Millipore Sigma) or 10 μM nocodazole (Millipore Sigma), we added the drug to the Lebovitz medium a minimum of 15 mins prior to imaging.

### Immunofluorescence: Cytoskeleton and Golgi Apparatus

We passaged HEK293T cells onto rectangular coverslips in 100 mm petri dishes ~24 hrs prior to fixation. This protocol was adapted from Jimenez et al. (55) and uses their fixation method. After fixation, we incubated the fixed cells with blocking buffer (2% BSA in PBS) for 30 mins with gentle rocking on an orbital shaker. Subsequently, we added a 1:400 solution of primary antibody in blocking buffer (anti-GM130 ab52649 Abcam, anti-tubulin Millipore Sigma T6199). We incubated coverslips with the primary antibody overnight at 4 °C. We then performed 3x 10-minute washes. After washing, we added a 1:400 dilution of fluorescent secondary antibody, goat anti-mouse 594 (A11032, ThermoFisher) and goat anti-rabbit (18772, Millipore Sigma) and incubated it for 1 hr at room temperature protected from light. We then washed as described after primary antibody incubation. To label actin, we added phalloidin (Millipore Sigma) labeled with Alexa-647 (0.094 μM in PBS) and incubated for 20 mins. We transferred coverslips to PBS, then sealed them to a glass slide for imaging. Finally, we imaged the cells in ~1.5-2 μm thick z-stacks using near-TIRF microscopy, with 1 mW laser power at 640 nm, 561 nm, and 488 nm excitation, 600 ms exposure time, EM gain of 200 using an EMCCD camera of a Nikon Eclipse Ti-E microscope.

### Immunofluorescence: Motor Colocalization

This protocol differs from cytoskeletal imaging in the fixation step; we incubated cells at −20 °C with 1 mM EGTA (BioShop) in −20 °C methanol, for 8 mins. Subsequent steps are identical to the cytoskeletal immunofluorescence protocol, except the blocking buffer contained 0.2% saponin (Millipore Sigma). We first imaged cells for the motor (kinesin or dynein) and saved their positions on the microscope. Following imaging, we washed the sample and probed for tubulin while maintaining the slide in the same position on the microscope, using another colour and shorter incubation times to allow higher throughput. The protocol was as follows: 1 hr incubation with primary antibody, wash 3x slide volume with blocking buffer, 45 mins incubation with secondary antibody, wash 3x slide volume with PBS, this sequential antibody staining was done due to the redundancy in antibody species available. The primary antibodies used for experiments were: 1/400 T9026 ms-alpha-tubulin, Millipore Sigma lot: 000089497, 1/400 ms-kinesin heavy chain (kinesin-1) MAB1614 lot: 3593684, 1/400 MAB1618 ms-dynein, cytoplasmic lot: 3089121, 1/400 ms-kif16b SAB1401759 lot: 09177-2B5, Millipore Sigma, 1/400 ms-kif1a cat:612094 lot: 9196885, BD Biosciences. Note this kif1a antibody has 47% sequence homology in the antibody binding site and 63% positives with the human kif1b sequence and therefore is likely to recognize both forms of kif1. 1/400 rb-kif3a ab11259 lot: BR183085-6, Abcam, this antibody could be used in conjunction with the tubulin antibody, rather than performing sequential staining. Secondary antibodies used for experiments were goat anti-mouse 488 (A21236, ThermoFisher Scientific) for motors followed by Goat anti-mouse 647 (A21236, Millipore Sigma) for microtubules.

### Immunofluorescence: Huntingtin Localization

We fixed the cells with prewarmed 4% paraformaldehyde (Millipore Sigma) at 37 °C for 10 mins, and subsequently followed the protocol as outlined for cytoskeletal imaging. The huntingtin antibody used was MAB2166 ms-huntingtin (Abcam lot: 3703568). The blocking buffer contained 0.2% saponin (Millipore Sigma).

### Immunofluorescence: Mitochondria

We fixed cells using prewarmed 4% paraformaldehyde (Millipore Sigma) at 37 °C for 20 mins, followed by 3 washes with PBS. Using 50 mM NH4Cl in PBS, we quenched fixation for 10 mins, and washed again 3 times with PBS. We permeabilized cells using 0.1% TritonX-100 in PBS for 10 mins with gentle rocking, followed by blocking with 10% BSA for 30-45 mins with gentle rocking. We diluted the primary antibody 1/500 (TOMM40 18409-1-AP, Proteintech, Rosemont, Illinois) with 5% BSA in PBS and incubated overnight at 4 °C. We washed coverslips 3 times with gentle rocking using the 5% BSA in PBS. We diluted the secondary antibody 1/400 in the same solution and incubated for 1 hr at room temperature. We repeated the washing described after the primary antibody incubation, then mounted the slide using PBS as the imaging medium. We imaged samples using an Olympus IX83 with a 100x objective using a disk scanning unit to obtain high resolution z stacking in a 25 μm range using 0.5 μm slices.

### Intracellular Optical Tweezers

We prepared a fibronectin (Corning, Bedford, Massachusetts) stock at a concentration of 1.4 mg/mL in water and stored it at −20 °C until use. We coated fluorescent yellow/green 500 nm beads (ThermoFisher Scientific) by centrifuging 20 μL of stock bead solution at 20000 rcf for 5 mins, then resuspended the beads in 25 μL of the fibronectin solution. We incubated the beads overnight at 4 °C, then washed with 40 μL PBS 3 times, centrifuging again at 20000 rcf for 5 mins each. We diluted this bead solution 1/20 in DMEM complete media, sonicated for 5 mins, and incubated with the cells for 10 mins at 37 °C. For imaging, we replaced the medium with Lebovitz supplemented with 10% FBS. Early endosome data was acquired from 15 mins–50 mins after bead internalization, while lysosomes were from 1 - 2 hours post-internalization. We identified cargoes moving processively within the cell, positioned the location of the center of the optical tweezer laser above the motile cargo, and opened the shutter to capture the bead. We acquired 60 s force traces holding the optical tweezer laser (12 W, 1064 nm) fixed in the original capture position, while monitoring the position signal and live image to ensure that the bead remained within one bead radius of the center of the tweezer. Immediately after this measurement, we calibrated the optical tweezers by applying a 275 s multifrequency sinusoid to calculate the position sensitivity (β), trap stiffness (ktrap), and viscoelastic properties (G’, G’’, **Figure S3e, f**) of the cytoplasm as previously described (56).

### Immunofluorescence Analysis: Actin, Microtubules, Golgi, Huntingtin

For each z-stack, we chose the sharpest image for each channel for qualitative analysis and observation as overlays.

### Immunofluorescence Analysis: Cell Morphology

We detected the cell edges from immunofluorescence images of actin using Cellpose software (57) with a cell diameter set to 225 pixels, flow threshold of 0.4, and using the “cyto” model. All partial cells at the edges of the field of view were manually removed from the analysis. This software generated regions of interest that we loaded into ImageJ and used to analyze the cell shape, area, and actin intensity.

### Mitochondrial Immunofluorescence Analysis

We determined the range of z-values with samples in focus, and only used those slices for image analysis. We processed the images in Fiji with the white top hat 3D morphological filter, using a radius of 5 in x and y, and 2 in z. We then set an automatic threshold and converted the thresholded image to a mask. Next, we ran the analyze particles function to determine the shape, area, and number of mitochondria in each plane, repeated for all in focus z planes, and added all the values for each z plane together. We plotted the results from all cells using MATLAB.

### Golgi Dispersal Analysis

Using ImageJ, we observed each of the Golgi images using the same linear LUT (minimum = 145 a.u. and maximum = 1885 a.u.) and then categorized the Golgi into organized, dispersed, or undetermined. Comparing the results from multiple experiments using this binary categorization and excluding the undetermined data, we determined the percentage of organized and dispersed Golgi observed in each cell type.

### Tracking Analysis

We used the Fiji plugin TrackMate (58) to generate tracks from the images for subsequent analysis. The settings used in TrackMate influence the results when they are inappropriately assigned, with parameters that are too stringent resulting in short tracks, and parameters that are not stringent enough resulting in linking of multiple tracks and signal noise. We approached choosing the parameters by comparing the maximum intensity projection (raw data) of all frames with the generated tracks, to ensure the generated tracks matched the raw data, and systematically attempting multiple values that would be considered logical for these cargos (**Tables 1 and 2**). The results from TrackMate are robust when the values are chosen within a logical range for a given single particle tracking experiment (58).

**Table 1.** TrackMate analysis settings used in the analysis of lysosomal cargoes using LysoTracker™.

| Parameter | Value |
| --- | --- |
| Estimated blob diameter | 1 $\mu\text{m}$ |
| Intensity threshold | 400 |
| Linking maximum distance | 0.5 $\mu\text{m}$ |
| Gap-closing maximum distance | 0.5 $\mu\text{m}$ |
| Gap-closing maximum frame gap | 2 |

**Table 2.** TrackMate analysis settings used in the analysis of early endosomal cargoes using Rab5-eGFP transfected cells.

| Parameter | Value |
| --- | --- |
| Estimated blob diameter | 0.8 $\mu\text{m}$ |
| Intensity threshold | 300 |
| Linking maximum distance | 0.5 $\mu\text{m}$ |
| Gap-closing maximum distance | 0.5 $\mu\text{m}$ |
| Gap-closing maximum frame gap | 2 |

### Motility Assay Analysis-Quantitation

To analyze the raw TrackMate data, we wrote custom MATLAB codes, available on GitHub: https://github.com/hendricks-lab.

We determined the directionality of trajectories by calculating the on- and off-axis displacements with respect to a line extending from the cell center to the cell periphery. Positive displacements correspond to outwards movements and negative displacements correspond to inwards movements. We next identified individual runs by finding reversals. Using mean-squared displacement analysis, we categorized processive runs as runs > 0.4 μm for early endosomes and >0.6 μm for lysosomes (**Figure S5g-j**). The processivity parameter α for processive runs should be greater than 1, α of diffusive runs should be ~1 as defined for Brownian motion, and there should be no directional bias for diffusive runs (**Figure S5g-j**). We then calculated the fraction of time of processive runs moving outwards and inwards. Diffusive runs were defined as any run between the minimum run length threshold and 0.03 μm, all runs below 0.03 μm were considered stationary.

We calculated time averaged mean squared displacement according to Eq. 1:

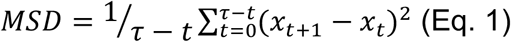

We calculated α values by fitting a line to the log-log plot of the mean squared displacement for time delays from 120 ms-5 s. We obtained radius of gyration values using Eq. 2:

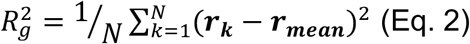

### Optical Tweezers: Force Trace Analysis

We calculated the trap stiffness and position sensitivity from the calibration data as previously described (56). Using the calibration data, we converted 60 s position traces to force traces (**Figure S2d**) and calculated the viscoelastic moduli in each condition (**Figure S2b, c**). We observed brightfield images of the position of the trap and the cell’s nucleus to define the directionality of the cargo’s movement. Any force event that had a maximum off-axis force, a force in the direction perpendicular to the defined microtubule axis, above 75% of the median on-axis forces, forces along the direction of the microtubule, were excluded. Stall forces were binned into 1 pN size bins and counted as stalls if the duration that they were above 0.5 pN duration was longer than 0.1 s. We calculated binding rates from the force traces by identifying the times that the cargoes unbound from the microtubules after a stall or unbinding event and determining the time between unbinding and rebinding to the microtubule. We calculated force-dependent unbinding rates from the stall force event before a cargo became detached and categorized them by the direction the cargo was travelling immediately prior to detachment, as previously described (59). This analysis assumes that the unbinding rate for these cargoes transported by multiple motors can be approximated by a single exponential (60). We performed an exponential fit to the data and presented it in Table S1 following Eq. 3 below with U as the unbinding rate, U_0_ as the force-independent unbinding rate, F as the force, and F_d_ as the detachment force.

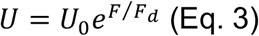

### Statistical Analysis

We analysed all pairwise histogram comparisons and cumulative distribution functions using the Kolmogorov-Smirnov (ks) test. We used Student’s two-tailed t-test for all pairwise analyses of violin plots. For optical tweezers stall force distribution analyses, in addition to the ks test we performed bootstrapping. We took 1000 bootstrap means of samples equivalent to the number of datapoints for the condition being tested. We then calculated 95% confidence intervals from the bootstrap mean distribution using the quantile function in MATLAB. To calculate p-values for bootstrap samples (not reported in **Figure 3**), we subtracted the control from the HTT-S421D condition for each of the 1000 bootstrap means for the conditions to be compared and plotted the distribution (**Figure S2e**). The p-value was determined by the fraction of data that overlapped with the 0 value, if this fraction was lower than 0.05, the difference between the values was considered statistically significant.

**Figure 2.**
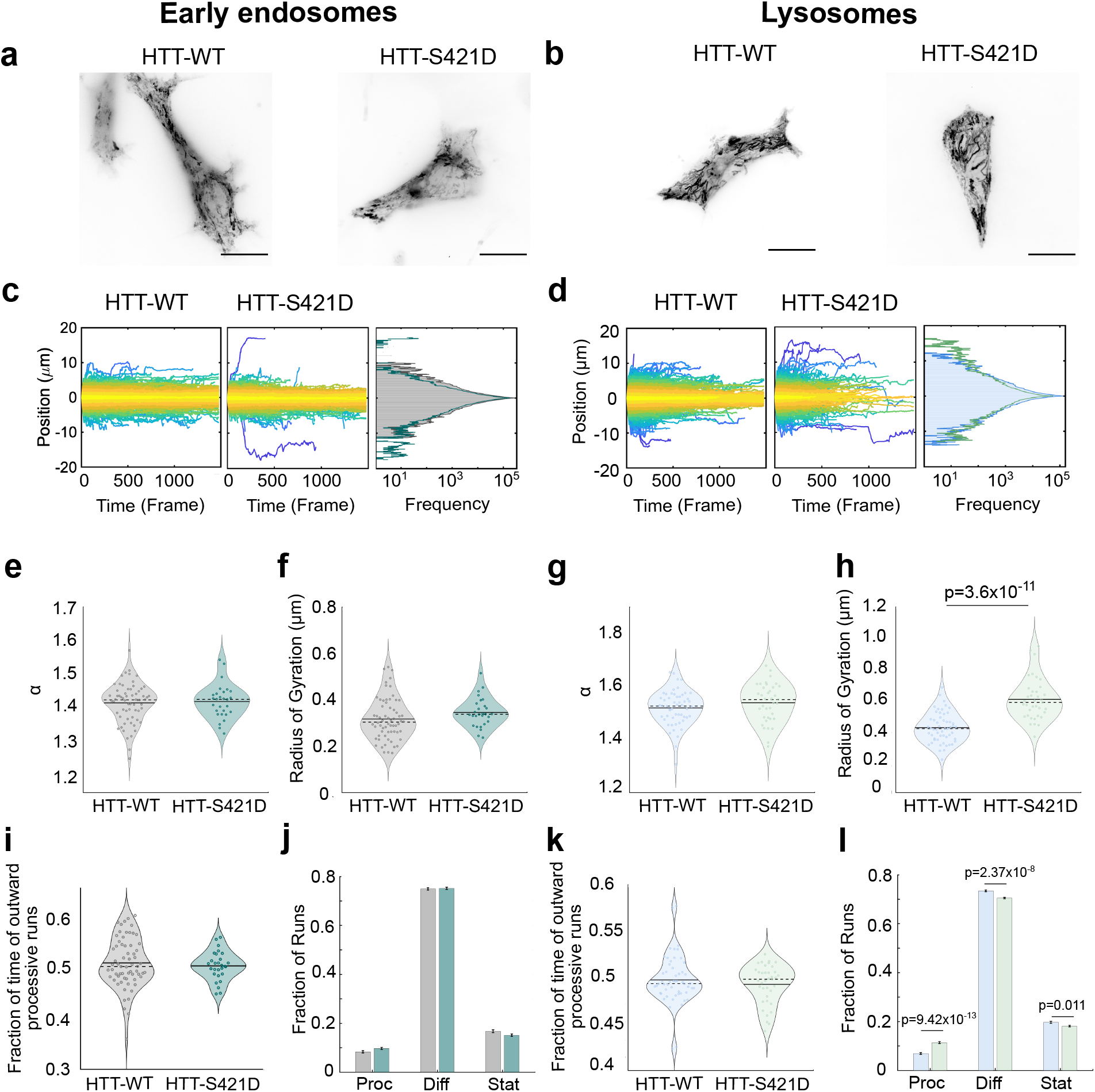
Huntingtin S421D mutation influences displacement and processive fraction of early endosomes and lysosomes. (**a, b**) Maximum intensity projection of early endosomes, labelled with Rab5-eGFP, and lysosomes, labelled with Lysotracker, for endogenous HTT-WT and HTT-S421D expressing cells. All scale bars are 20 μm. (**c, d**) All trajectories for early endosomes (**c**) and lysosomes (**d**) in both HTT-WT and HTT-S421D cells, coloured by their displacement (yellow indicates stationary while purple indicates long displacement) with their inward (in) motility indicated by negative values and outward (out) directionality indicated by positive values. Distributions of all positions are indicated on the left, with the early endosome HTT-WT in grey and HTT-S421D in teal, and the lysosome HTT-WT in blue and HTT-S421D in green. (**e, g**) Early endosome (**e**) and lysosome (**g**) processivity per cell, measured by α, the slope of the log-log plot of their mean-squared displacement. Means are represented with a filled line, while medians are dashed lines. (**f, h**) Radius of gyration (Rg) per cell of early endosomes (**f**) and lysosomes (**h**). Rg is a measure of the radius that contains half of the datapoints in the trajectory centered around its average position. (**i, k**) Early endosome (**i**) and lysosome (**k**) directional bias per cell of processive trajectories toward plus ends of microtubules (outwards), the remaining fraction of processive trajectories move inwards. Directional bias is determined by fraction of time particles are moving outwards compared to inwards and averaged for each cell. (**j, l**) The fraction of total runs within a trajectory categorized as processive (Proc), Diffusive (Diff), and Stationary (Stat) for early endosomes (**j**) and lysosomes (**l**). Colour codes match that in the previous figures with HTT-WT in grey for early endosomes and blue for lysosomes and HTT-S421D in teal for early endosomes and green for lysosomes. Error bars indicate standard error of the mean. The number of experiments, cells, and trajectories for each condition are the following; HTT-WT early endosomes: 65 cells, 41994 trajectories, 6 experiments; HTT-S421D early endosomes: 28 cells, 25476 trajectories, 3 experiments; HTT-WT lysosomes: 55 cells, 25247 trajectories, 5 experiments; HTT-S421D lysosomes: 38 cells, 12670 trajectories, 4 experiments; Statistical significance was determined via Student’s two-tailed t-test and p-values are indicated on each plot where applicable.

**Figure 3.**
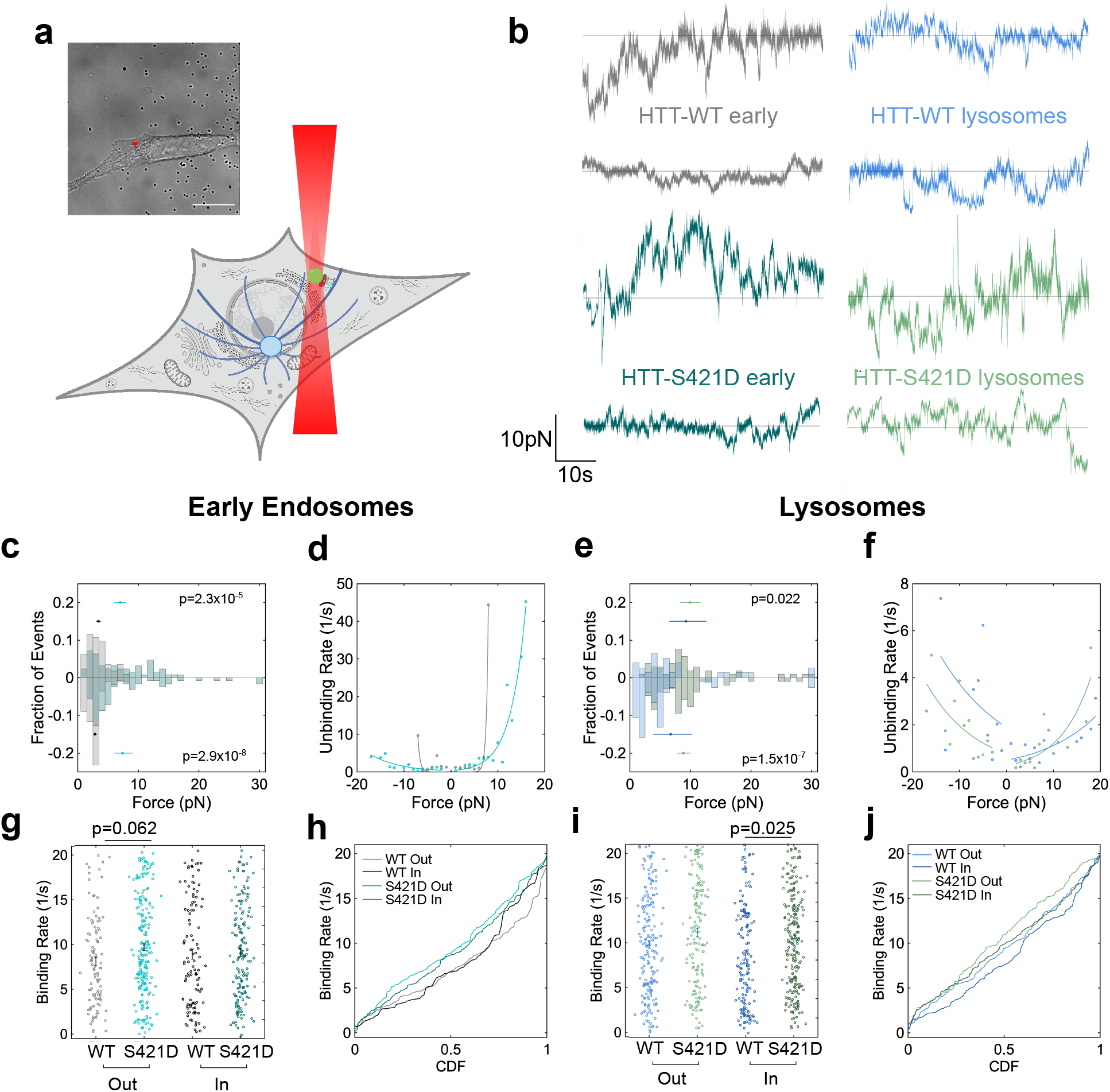
Force trace analysis demonstrates HTT-S421D induces higher binding rates, stronger binding, and higher forces in both directions for early endosomes and lysosomes. (**a**) Schematic of the optical tweezers assay including the endocytosed bead (green) inside the optical trap (red) along with a brightfield image of a cell during the experiment (top right). The scale bar is 20 μm and the arrowhead indicates the position of the optical trap during the experiment. (**b**) Sample force traces for early endosomes (early) and lysosomes in HTT-WT and HTT-S421D cells. The horizontal line indicates the zero axis, while positive values indicate outward forces and negative values indicate inward forces. (**c, e**) The fraction of early endosome or lysosome stall events at a given force with a minimum stall duration of 0.1 s, separated into 1 pN bins with the mean and 95% confidence intervals of bootstrap sampled means and p-values for each condition are shown above (for outward) or below (for inward) the distribution. The colours in (**c-j**) follow the colour legend from (**b**). (**d, f**) Force-dependent unbinding rates in both outward and inward directions (filled circles) for early endosomes or lysosomes fit with a single exponential (solid line). (**g-j**) Binding rates for each stall event in early endosomes and lysosomes of HTT-WT and HTT-S421D in both directions in (**g, i**) including the cumulative distribution function (CDF) for each condition in (**h, j**). Statistical significance by ks test is indicated by the p-value in the figure panel where applicable. All results from optical tweezers data for HTT-WT early endosomes from 10 cells, 5 experiments; HTT-S421D early endosomes: 16 cells, 8 experiments; HTT-WT lysosomes: 12 cells, 5 experiments, HTT-S421D lysosomes: 11 cells, 4 experiments. Statistical significance was determined by ks test for binding rates and force distributions and p-values are indicated where applicable.

## RESULTS

### Endogenous HTT-S421D mutation does not affect cytoskeletal organization or huntingtin localization

In generating the HTT-S421D HEK293T cell line, we aimed to preserve the native expression levels and stoichiometry of interactions between HTT and its many interactors. We used a lentiviral CRISPR cassette to generate this S421D point mutant cell line (**Figure 1b**) and identified a single positive clone with the desired mutation out of 53 monoclonal cell lines from puromycin selection (**Figure 1c**). We verified that cell ploidy remained the same after CRISPR editing (**Figure S1a-c**). HTT-S421D did not result in appreciable changes to the actin or microtubule organization, nor overall cellular morphology (**Figure 1d, Figure S1e, f**). However, actin intensity is higher in HTT-WT cells compared to HTT-S421D, suggesting HTT-S421D may lower F-actin concentration or bundling (**Figure S1d**). To determine whether HTT-S421D caused major defects in dynein function, we probed the organization of the Golgi apparatus (33, 61, 62). We observed no changes in the percentage of cells with organized versus dispersed Golgi, indicating dynein function was not severely disrupted (**Figure 1e**). Though there are potential off-target effects from any CRISPR gene editing (63), we have determined no major morphological, motor protein, or cytoskeletal defects. Importantly, the HTT-S421D mutation did not affect huntingtin’s localization nor expression level (**Figure 1f, g, Figure S2**). Overexpression of mCherry-HTT-WT (WT+HTT-WT) or mCherry-HTT-S421D (WT+HTT-S421D) leads to elevated expression of both constructs at similar levels (**Figure 1g**). In iPSC-derived neurons expressing polyglutamine-expanded huntingtin (180Q), HTT-S421D led to increased mitochondrial surface area while the number of mitochondria remained similar (44). Immunofluorescence of the mitochondria in HEK293T cells showed a decrease in the mitochondrial number and area upon HTT-S421D mutation (**Figure 1h-j**), suggesting that HTT-S421D mitochondrial regulation may be cell-type dependent. These results are not consistent with the previous study, however, results from the previous study are not directly comparable since they are derived from a disease cell line of a neuronal cell type and the mitochondria were analyzed in 3D compared to the current 2D analysis. Together, immunofluorescence and huntingtin expression level characterization demonstrated that the HTT-S421D mutation did not substantially alter the cytoskeleton, Golgi apparatus, nor localization of huntingtin while mitochondrial morphology was affected.

### Early endosome motility is weakly enhanced by huntingtin phosphorylation

We hypothesized that the processivity, displacement, and fraction of outwardly directed early endosomes would increase upon S421 phosphorylation based on previous work with BDNF vesicles in neurons (39). We transfected cells with Rab5-eGFP to fluorescently label early endosomes (**Figure 2a, c** and **Movies 1, 2**). We quantified the processivity of organelle trajectories using the slope of the log-log plot of mean-squared displacement (α). For cargoes transported by motor proteins, α >1 is indicative of directed transport while α = 1 represents diffusive motion. The radius of gyration (Rg) is an indicator of displacement that corresponds to a radius containing half of the points in the trajectory from its average position. To evaluate the directional bias and fraction of processive, diffusive, and stationary runs, a minimum run length for processive runs was set at 0.4 μm for early endosomes. At this threshold, the diffusive runs had an α value of ~1 and no directional bias (0.5 fraction outwards and inwards), suggesting the runs are purely diffusive (**Figure S5g, i**). In addition, the processive runs at the set minimum run length threshold have an α value greater than 1 to indicate their motion is processive (**Figure S5g**). The endogenous HTT-S421D mutation qualitatively resulted in a small but insignificant increase in the displacement of early endosomes (p=0.14 by two-tailed t test) without changing their overall processivity or directional bias (**Figure 2e, f, i**). However, the fraction of processive runs also showed a small but insignificant increase in the HTT-S421D cells (p=0.072 by two-tailed t test), while the diffusive and stationary runs remained similar between HTT-WT and HTT-S421D cells (**Figure 2j**). We expected an outward directional bias as previously described when truncated HTT-WT and HTT-S421D were overexpressed in primary neurons (23, 39). The lack of outward bias observed may be dependent on cell type, as neurons are highly polarized cells with specialized motors to modulate tight control of cargo directionality. This indicates that S421 phosphorylation is weakly increasing motor affinity for and activity on the microtubule or recruitment to early endosomes, with no preference for kinesins or dyneins.

To dissect the effects of HTT phosphorylation and expression levels, we transiently transfected full length mCherry-HTT-WT (WT+HTT-WT) or mCherry-HTT-S421D (WT+HTT-S421D). These transfections lead to a ~4-10-fold increase in HTT expression (**Figure 1g**). Overexpression of WT+HTT-S421D increased the processivity and displacement of early endosomes without impacting their directionality (**Figure S3a, b, e**). HTT-S421D overexpression increased the fraction of processive runs and decreased the fraction of diffusive runs of early endosomes (**Figure S3f**). These unexpected results in overexpression conditions may be due to the change in stoichiometry of HTT’s interactions with its native binding partners. The gene edited HTT-S421D HEK293T cells demonstrate a slight increase in processive fraction of early endosomes, while HTT overexpression alters motility.

**Movie 1. Live cell sample video of wild-type Rab5-eGFP early endosomes used for tracking analysis.** Image colours are inverted, the time scale is in seconds and frame rate is 10 fps, the scale bar is 20 μm.

**Movie 2. Live cell sample video of HTT-S421D Rab5-eGFP early endosomes used for tracking analysis.** Image colours are inverted, the time scale is in seconds and frame rate is 10 fps, the scale bar is 20 μm.

### HTT-S421D increases displacement and processive fraction of lysosomes

We predicted HTT-S421D’s effects would be universal for all HTT-associated cargoes, and therefore expected increased outward bias, processivity, and displacement of lysosomes. This hypothesis is supported by previous work showing that HTT promotes outward motility in the mid-axon of neurons for autophagosomes, which eventually mature into lysosomes (18, 19). In both HTT-WT and HTT-S421D experiments, the raw trajectories demonstrated bidirectional motility (**Figure 2b, d** and **Movies 3, 4**). We evaluated the minimum run length for processive runs in lysosomes the same way we did for early endosomes and found the threshold that met the defined criteria to be 0.6 μm (**Figure S5h, j**). HTT-S421D induced a significant increase in displacement, without significantly affecting their processivity (p=0.19 by two tailed t-test) or directional bias (p=0.36 by two tailed t-test), although both qualitatively appear mildly increased (**Figure 2g, h, k**). The fraction of processive runs is significantly higher in cells expressing HTT-S421D compared to HTT-WT, while the diffusive and stationary fractions are lower (**Figure 2l**). When we overexpressed HTT, we observed a minor increase in processivity for HTT-WT and a substantial increase in displacement regardless of the phosphorylation state (**Figure S3d, e, f**). Unexpectedly, HTT-WT and HTT-S421D overexpression led to an inward directional bias (**Figure S3g**). Interestingly, the fraction of lysosome processive and stationary runs are higher when HTT is overexpressed, while the diffusive runs decrease (**Figure S3h**). Taken together, our results indicate that HTT overexpression modifies processivity, displacement, and directional bias while phosphorylation increases their displacement and fraction of processive runs. This suggests S421 phosphorylation at native expression levels is increasing the binding rate, recruitment, or activity of lysosome-associated kinesin and dynein motors more significantly than for early endosomes.

**Movie 3. Live cell sample video of wild-type LysoTracker™ lysosomes used for tracking analysis.** Image colours are inverted, the time scale is in seconds and frame rate is 10 fps, the scale bar is 20 μm.

**Movie 4. Live cell sample video of HTT-S421D LysoTracker™ lysosomes used for tracking analysis.** Image colours are inverted, the time scale is in seconds and frame rate is 10 fps, the scale bar is 20 μm.

### Tethering on actin limits the motility of early endosomes

Huntingtin interacts with optineurin, which recruits myosin VI and regulates cargo association to actin (32). We used latrunculin A (latA) to partially depolymerize actin filaments and reduce the contribution of actin-based motility and tethering. We expected an increase in displacement and processivity upon latA treatment since cargo tethering and confinement by actin are reduced, allowing long range, processive motility by microtubule motors to dominate. We determined the optimal concentration for motility assays via imaging cells labelled with SiR-actin. We determined 250 nM latA was optimal as it resulted in significant depolymerization of actin filaments without loss of native cellular morphology (**Figure S4a**). As expected based on previous work (64), treatment with latA increased displacement and processivity of Rab5 early endosomes (**Figure S4b, c**). HTT-WT expressing cells showed no change in directional bias upon latA treatment, indicating that allowing both kinesin and dynein to dominate transport does not impact its direction (**Figure S4d**). HTT-S421D early endosomes no longer maintain the slight increase in directional bias upon latA treatment, indicating that these cargoes may already be localized to microtubules in the native condition. This may be due to an inherently lower concentration of F-actin observed in the HTT-S421D cells, which could reduce cargo tethering and increase their microtubule-based transport (**Figure S1d).** The increased processivity and displacement in the latA condition is similar to our observations in **Figure 2,** suggesting that both loss of actin tethering and HTT phosphorylation act through similar mechanisms.

### Microtubule-based transport dominates lysosome motility

We used 10 μM nocodazole (NZ) to depolymerize microtubules and observe the magnitude of the impact on the motility parameters. We expected that upon NZ treatment, both processivity and displacement would decrease because microtubule-based motors drive the long range processive transport of both early endosomes and lysosomes. We observed this expected decrease in displacement and processivity for lysosomes, suggesting that they rely heavily on microtubule-based transport to drive their motility (**Figure S4f, i, j**). The substantial change in processivity and displacement for lysosomes demonstrates the magnitude of changes in these parameters upon large scale disruption of the cytoskeleton. Early endosomes demonstrated a surprising lack of effect of nocodazole treatment on their processivity and displacement, perhaps due to their inherently low processivity and displacements compared to lysosomes (**Figure S4e, g, h**). This data contrasts with a previous study which showed early endosome transport is affected by nocodazole, however they used a higher concentration of nocodazole, and they labeled cargoes with quantum dots which may include later stages of cargoes in the endosomal pathway (64). The lysosomes in NZ-treated cells reached similar values for processivity and displacement to the early endosomes in the control condition, suggesting that the values for early endosome motility were close to their minimum. Combining this result with our conclusions from the latA treatment demonstrates that early endosome transport may be more dependent on switching between actin-based and microtubule-based transport (**Figure S4**). These results indicate early endosome transport has a lower dependence on microtubules compared to lysosomes, perhaps due to its inherently short displacements and lower processivity.

### S421 huntingtin phosphorylation activates kinesins and dyneins on early endosomes

To determine how changes in motor activity contribute to the motility we observed in **Figure 2**, we measured the force generation of motors on early endosomes with optical tweezers. We incubated cells with fibronectin-coated 500 nm polystyrene beads for 10-50 mins (65) and used optical tweezers to measure the forces that motors exerted in living cells (**Figure 3a, b**). At this stage of maturation, the early endosomes are positive for Rab5 and other early endosome markers and represent a subpopulation of the vesicles labeled by Rab5-eGFP. We analyzed cargoes that were moving processively and bidirectionally to obtain measurements for early endosomes most likely being transported along microtubules, however it is possible some actin-based motor activity contributed to the forces measured (66). During calibration, we noted that the material properties of the cytoplasm for HTT-WT and HTT-S421D conditions were not significantly different, as expected (**Figure S6 a, b**). Early endosome force traces demonstrated the expected bidirectional motility biased inwards (**Figure 3b, Figure S6d**). We predicted the HTT-S421D mutation might weakly increase motor activity on early endosomes, given the weak increase in displacement and processive fraction (**Figure 2f, j**). Interestingly, the stall force distributions for early endosomes shifted towards higher forces in both outward and inward directions with HTT-S421D (**Figure 3c**), suggesting both kinesins and dyneins were activated. In cells, each kinesin motor exerts forces of 2-7 pN, and each dynein exerts approximately 2-3 pN (67). Looking closely at the distributions, we observe frequent kinesin forces around 3 pN and 6 pN for cells with HTT-WT (**Figure 3c**). Both of those populations are observed in HTT-S421D cells, in addition to high force events near 9 pN and 12 pN. Observing the inward directed forces, HTT-WT cells show frequent dynein forces near 1.5 pN and 5 pN, while forces shift to ~ 3.5 pN, 9 pN, and 13 pN in HTT-S421D cells. Together, we estimate based on the literature that these forces likely correspond to ~1-2 active kinesins and 1-3 active dyneins in HTT-WT cells, and 1-4 active kinesins and 2-6 active dyneins in HTT-S421D cells, which is like previous estimates for late phagosomes (68, 69). The outward forces are not significantly different from the inward forces in both conditions, which we expected based on the results from **Figure 2**. This suggests that the kinesins are being activated to the same level as the dyneins, increasing the cargo’s engagement with the microtubule.

We then sought to determine other motor-relevant parameters to determine how the force-dependent unbinding rates of each motor were affected by HTT-S421D, predicting that the kinesin and dynein motors would have a higher force-dependent resistance to load, therefore inducing the higher processive fraction (**Figure 2j**). The unbinding rates of kinesin and dynein teams on early phagosomes in HTT-WT cells is similar (**Figure 3d, Table S1**). We observed slower unbinding under load of both kinesin and dynein teams with HTT-S421D, consistent with more kinesin and dynein motors engaged with the microtubule (**Figure 3d, Table S1**). Next, we measured the binding rate of early endosomes moving inwards and outwards and found the binding rate for early endosomes was increased for both directions in HTT-S421D cells (**Figure 3g, h**). The binding rate was more strongly enhanced for outward forces, suggesting an increased number of active kinesins enhances binding compared to HTT-WT (**Figure 3g**). Observing the cumulative distribution functions, we note that outward and inward motor binding rates in HTT-S421D cells were similar (**Figure 3h**). In summary, the force data indicate that huntingtin phosphorylation at site S421 increases the net force generation, affinity, and microtubule binding strength of both kinesin and dynein motors on early endosomes.

### HTT-S421D increases lysosome-associated kinesin and dynein engagement with microtubules

Like for early endosomes, we hypothesized that kinesins would have increased activity on lysosomes as they also demonstrated an increased displacement, processive fraction, and slight increase in outward motility, indicating kinesin activity may be favoured (**Figure 2h, k, l**). We measured the force characteristics of phagolysosomes by modifying the optical tweezers assay described for early endosomes with a longer incubation time of 1-2 hrs. The phagolysosomes measured in the optical tweezer experiments, termed lysosomes hereafter, represent a subpopulation of lysotracker-positive lysosomes, compared to our earlier analysis tracking all lysotracker-positive vesicles (**Figure 2**). Sample force traces demonstrated typical bidirectional motility of lysosomes (**Figure 3b, Figure S6d**). We observed an increase in forces in both directions for lysosomes as for the early endosomes, although to a lesser degree (**Figure 3e**). The balance of kinesin to dynein forces remained similar in both control and HTT-S421D conditions, as expected for these bidirectional cargoes (**Figure S6e**). Predicting the approximate number of motors as described for early endosomes, the force distributions in the HTT-WT condition contain ~1-3 active kinesins and ~1-2 active dyneins, while the HTT-S421D condition displays ~1-2 additional kinesins and ~1-2 additional dyneins are engaged. Like the results in early endosomes, the outward forces were not significantly higher than the inward forces under any condition, which indicates the forces alone do not determine the overall directional bias. The unbinding rates indicate that kinesin teams are more resistant to unbinding than dyneins for lysosomes (**Figure 3f**). In HTT-S421D cells, we observed a slightly lower resistance to unbinding for lysosomes moving outward, while the unbinding resistance was higher than HTT-WT for inward-moving cargoes (**Figure 3f, Table S1**). Finally, we examined the binding rates for lysosome motors to determine how HTT-S421D affected microtubule affinity. HTT-S421D increased the binding rate for both inward- and outward-directed cargoes compared to HTT-WT (**Figure 3i, j**). The increased forces and enhanced binding we observed in HTT-S421D cells suggests HTT phosphorylation recruits and/or activates both kinesins and dyneins.

### HTT-S421D increases kinesin-1 and dynein association to microtubules

Since we observed increased activity of both kinesin and dynein motors on early endosomes and lysosomes upon S421 phosphorylation, we asked if more kinesins and dyneins were recruited to microtubules to generate the observed increased displacement and processive fraction of early endosomes and more significantly, lysosomes. Previous studies overexpressing HTT-S421D demonstrated that kinesin-1 increases its association with microtubules (39). We used immunofluorescence imaging to approximate the differences in the number of motors associated with microtubules in each cell type. We fixed the cells with methanol to remove cytosolic components and determine the relative number of motors that were bound to microtubules in each condition at the time of fixation (**Figure 4a, b**). We then created intensity heatmaps of all images from the microtubules and kinesin-1 heavy chain (KHC) or dynein intermediate chain (DIC) and plotted them on the same axes (**Figure 4c-f**). We detected higher intensities in the motors channel for both KHC and DIC in the HTT-S421D cells (**Figure 4c-f**). The higher intensity indicates a higher concentration of motors on extracted microtubules in the HTT-S421D cells. Upon calculating the correlation coefficients between the microtubule and motor channels for each individual image, we observed that the dynein and microtubule intensities were more highly correlated in HTT-S421D cells than WT cells. This assay indicates that HTT-S421D increases the fraction of kinesin-1 and dynein motors associated to microtubules. To determine if other kinesins are affected by HTT-S421D, we compared the correlation of kif3a (kinesin-2), kif1, and kif16b (both kinesin-3) on microtubules. We observed that the HTT-S421D mutation decreases the amount of kif3a and kif16b on microtubules compared to HTT-WT, suggesting kinesin-1 is the dominant plus-ended motor in cargo complexes with phosphorylated huntingtin (**Figure S7**). These results support our data from the optical tweezers analysis, which showed a higher binding rate and lower force-dependent unbinding rate for motors in the HTT-S421D cells in both early endosomes and lysosomes. Further, the signal for microtubule-bound kinesin-1 and dynein motors is higher in HTT-S421D cells, suggesting HTT S421 phosphorylation increases the engagement of both kinesins and dyneins with microtubules.

**Figure 4.**
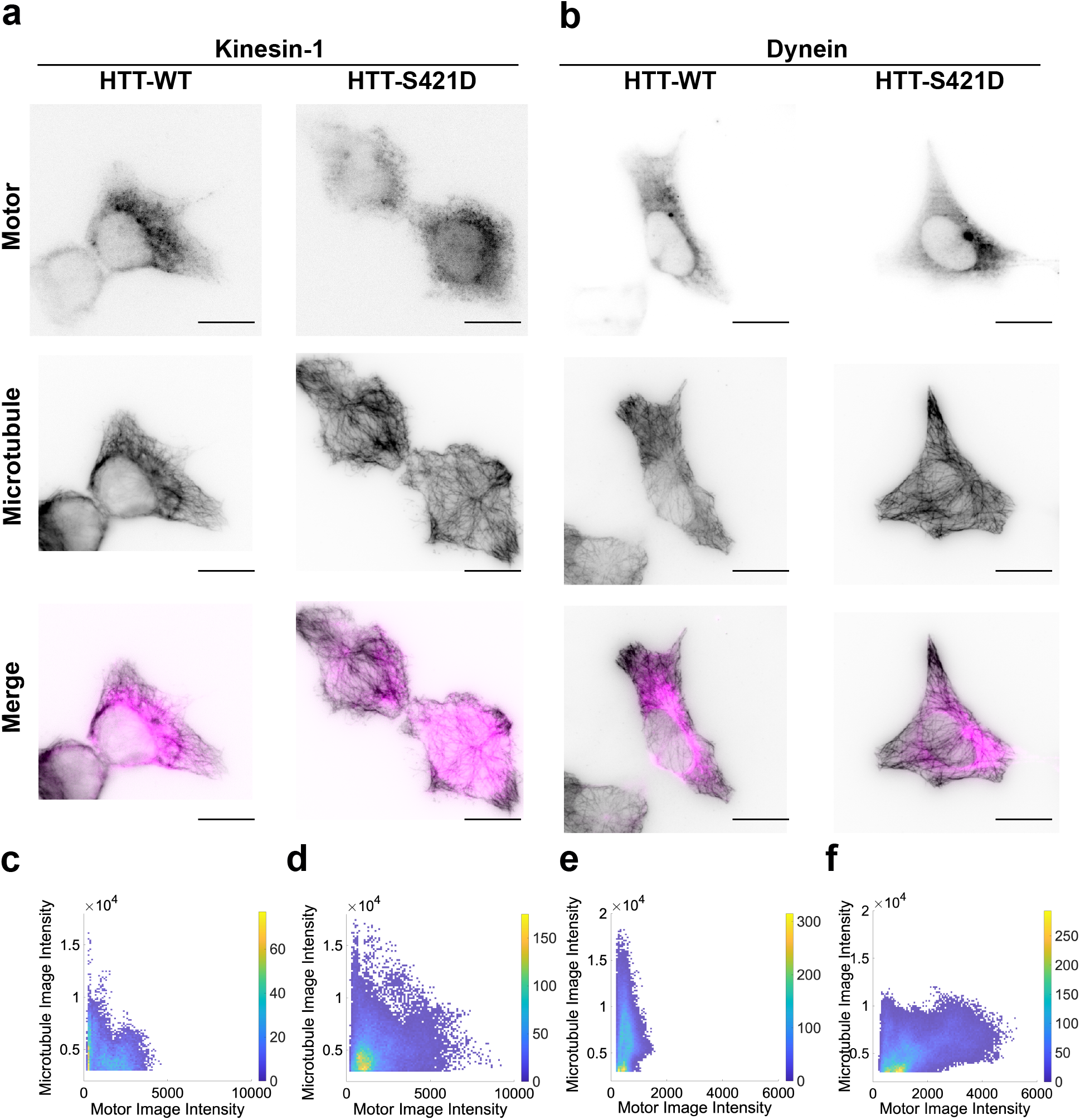
HTT-S421D increases kinesin and dynein association to extracted microtubules. (**a, b**) Inverted immunofluorescence images demonstrating colocalization (bottom) kinesin heavy chain (kinesin-1, (**a**), top) or dynein intermediate chain (dynein, (**b**), top) on extracted microtubules (middle) in HTT-WT and HTT-S421D huntingtin cells. In the merge channel, motors (kinesin-1 or dynein) are indicated in magenta and microtubules in black. All scale bars are 20 μm. (**c-f**) Colocalization plots were created from plotting intensity values of each pixel of the microtubule and motor (kinesin or dynein) channels of all images. Differences between kinesin-1 HTT-WT and HTT-S421D correlation coefficients resulted in p=0.4748, while dynein HTT-WT compared to HTT-S421D correlation gave p=0.0533 by a two-tailed Student’s t-test. Data for HTT-WT kinesin-1 from 17 cells, 3 experiments, HTT-S421D kinesin-1: 15 cells, 2 experiment, HTT-WT dynein: 32 cells, 4 experiments, and HTT-S421D dynein: 32 cells, 4 experiments.

## DISCUSSION

Our results show that huntingtin phosphorylation at S421 increases the number of active kinesin and dynein motors on endosomes. Surprisingly, huntingtin activates microtubule motors on endosomes at varying stages of maturation, from early endosomes to lysosomes. Further, huntingtin enhances the force generation of both kinesin and dynein teams in living cells. Huntingtin also regulates the motility of neuronal signalling vesicles including BDNF and APP (22–24, 39). In contrast, most cargo adaptors are recruited to specific cargoes at specific stages of maturation (19, 70). The ubiquitous function of huntingtin on a wide array of cargoes illustrates its role as a central regulator of vesicular transport.

While kinesin and dynein teams exert more force and have enhanced microtubule binding on both early endosomes and lysosomes in HTT-S421D cells (**Figure 3,4**), its effect on the motility of different endocytic cargoes varies. For early endosomes, huntingtin phosphorylation does not strongly affect motility (**Figure 2e, f**). However, the force generation and motor binding rates are significantly increased (**Figure 3**). Previous studies indicate that huntingtin’s effect on Rab5 cargoes is strengthened when cells were under stress (71, 72). Akt, the kinase responsible for phosphorylating HTT at S421, is activated upon cellular stress (73, 74). Therefore, the enhanced force generation by kinesin and dynein upon HTT-S421 phosphorylation may be a response to cellular stress, and we would expect that HTT phosphorylation might have a stronger effect on early endosome motility in stressed conditions. In comparison, lysosomal displacement and processive fraction is strongly enhanced in HTT-S421D cells (**Figure 2h, l**). The stronger effect of HTT S421 phosphorylation on lysosomes may be due to the consistent association between huntingtin and Rab7 cargoes, typically categorized as late endosomes or lysosomes, even in the absence of cellular stress (75).

Taken together, these results suggest that huntingtin phosphorylation regulates transport by enhancing the recruitment and activity of both kinesin-1 and dynein (**Figure 5a, b**). However, its effects on the motility of different cargoes depend on the identity and motility characteristics motility characteristics of the cargo in the control condition. Initially, early endosome motility consists of short displacements punctuated by pauses, while activating both kinesin and dynein by HTT phosphorylation results in an increased engagement with the microtubule and a slight increase in average displacement and processive fraction. Lysosome motility is equally distributed towards the cell center and periphery, such that activating kinesin and dynein has a weak effect on directionality, but further increases the total displacement and processive fraction of cargoes.

**Figure 5.**
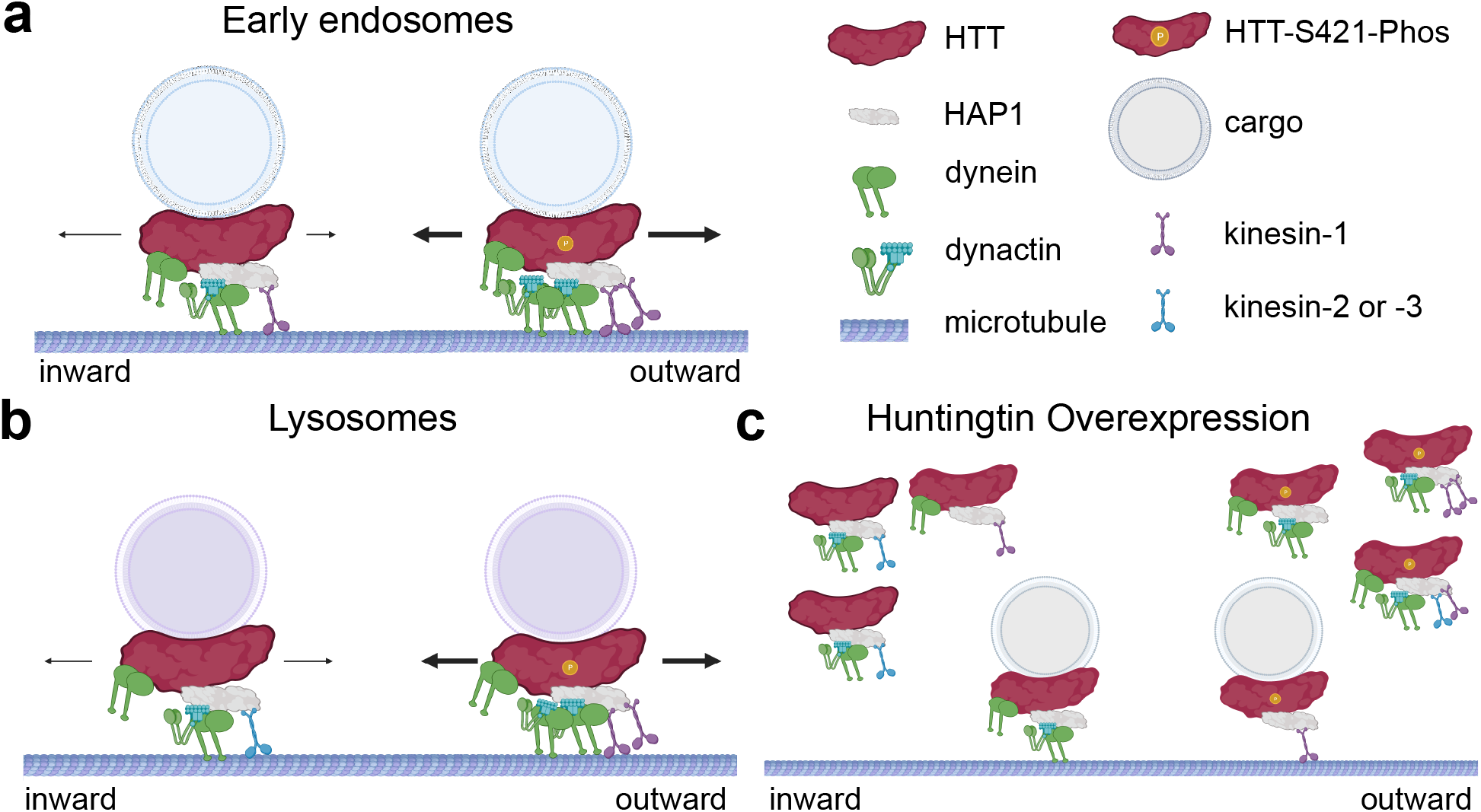
Model of huntingtin S421 phosphorylation and expression level mediating regulation of early endosome and lysosome transport. (**a**) Huntingtin S421 phosphorylation (HTT-S421-Phos) increases activity of kinesin and dynein motor proteins on early endosomes. (**b**) Lysosomes move bidirectionally. When kinesin and dynein activity are enhanced by huntingtin phosphorylation, the movement remains bidirectional, but the displacement and processive fraction of trajectories increases. (**c**) Overexpression sequesters motors and adaptors or disrupts vesicle-bound huntingtin-adaptor-motor complexes regardless of cargo identity or huntingtin phosphorylation state.

Increased displacement and processivity could result from either engaging more kinesin and dynein motors or by reducing tethering on actin. We observed that disrupting actin filaments resulted in an increased processivity and displacement of early endosomes similar to the effect of HTT-S421D on lysosomes (**Figure S4a-d**), and the actin intensity was reduced in HTT-S421D cells (**Figure S1d**), suggesting that reduced myosin VI-mediated tethering to actin filaments (32) could contribute to enhanced transport in the HTT-S421D cells (66). However, we also find that more kinesins and dyneins are bound to microtubules in HTT-S421D cells (**Figure 4)**. Further, optical trapping experiments showed that HTT-S421D results in enhanced forces for both kinesin and dynein (**Figure 3**). These results suggest HTT-S421 phosphorylation enhances transport through both reduced actin-mediated tethering and activating kinesin and dynein.

Our results support previous findings that overexpression of phosphomimetic HTT-S421D results in increased outward motility of BDNF and TI-VAMP vesicles compared to phosphoresistive HTT-S421A (23, 39), and knock-down and rescue with polyQHTT-S421D results in an outward bias of BDNF and APP vesicles compared to polyQ-HTT (22, 24). However, while previous studies observed increased kinesin-1 recruitment to microtubules and enhanced interactions between the p150glued subunit of dynactin and KHC (Kinesin 1 Heavy Chain) upon HTT-S421D expression (40) and decreased recruitment of kinesin-1 to APP vesicles upon expression of HTT-S421A (24), we observed an increase in recruitment of both motors. These differences between previous results and our observations may reflect effects of HTT expression levels. Alternatively, they may be due to differences in HTT function in the embryonic kidney cells used here compared to previous studies in neurons, which are highly polarized and have higher endogenous HTT expression. An important limitation of both previous works and this study is that the cell signalling state can be impacted by pseudo phosphorylation of this type and therefore could lead to secondary effects that may impact transport (76). In-depth characterization of cell signalling states would be an important future work to understand how pseudo phosphorylation of this ubiquitous protein affects other cellular processes. Previous studies were performed using overexpression in neurons, which have a unique cytoskeletal organization and express specific isoforms of motor proteins and tubulin to transport cargoes long distances along the axon. Current optical trapping methods are challenging to perform in neurons as the conventionally used beads for optical tweezers are too large to be endocytosed, and smaller beads of the same material do not have a high enough refractive index to remain in the trap. This study opens the door to combining advances in optical trapping methods with gene editing in pluripotent stem cell derived neurons to enable similar biophysical studies in neuronal systems in the future.

Overexpression of HTT alters the processivity, displacement, and directional bias of early endosomes and lysosomes. The effects of overexpression are highly variable and often do not reflect the trends observed when inducing mutations in the endogenous HTT (**Figure 2**), suggesting that maintaining the stoichiometry of HTT to its many interacting proteins is critical. We expect that excess HTT in the cytoplasm is interacting with and sequestering the motors and adaptors required to generate the effect that we observe in transport upon overexpression (**Figure 5c**). This result supports the known consequences of overexpression experiments, which cause adverse effects by changing the stoichiometry of interactions with other proteins in the cell. For example, overexpressing the p50 dynamitin subunit of dynactin causes a dominant negative effect in which early endosomes and lysosomes become peripherally localized due to dynein sequestration (62). Regardless of the phosphorylation state, overexpression of HTT affected the directional bias of early endosomes and lysosomes. Processivity and displacements were increased for lysosomes in cells overexpressing either HTT-WT or HTT-S421D, however, early endosomes only demonstrated this effect overexpressing HTT-S421D. This work, supported by many previous studies discussing expression levels (e.g. (62, 77–79)) highlights the importance of examining protein function at endogenous expression levels to ensure biological relevance of the effects. This is particularly important for scaffolding proteins like huntingtin, which mediates interactions among many adaptors, effectors, motors, and membranes.

Transport is tightly regulated on multiple levels within the cell to ensure specific cargoes arrive at their intracellular destination. Scaffolding proteins such as HTT are a primary mechanism of regulation that mediate interactions between cargoes and motor proteins, and are the only known level of regulation that is cargo specific (19, 39, 45). Many scaffolds specifically interact with a limited set of cargoes and are regulated by post-translational modifications (19, 45, 48, 80, 81). Here, we illustrate the role of HTT in regulating early endosome and lysosome transport. HTT S421 phosphorylation activates kinesin and dynein on both early endosomes and lysosomes. However, enhanced kinesin and dynein activity differentially affects early endosomes and lysosomes, influencing their direction, motor activity, and affinity for microtubules. Our results underscore the significant role of huntingtin in directing endosome transport and suggest that huntingtin mutations likely result in cargo-specific disruptions to both signalling and degradative pathways.

## AUTHOR CONTRIBUTIONS

E. N. P. P., A. R. C., and A. G. H. designed the study. A.R.C. and D. S. generated the CRISPR cell line. A.R.C. developed the codes for analysis. E. N. P. P. performed the experiments and analyses. E. N. P. P. and A.G.H. wrote the manuscript.

## DECLARATION OF INTERESTS

The authors report no competing interests.

## ACKNOWLEDGEMENTS

We thank Frederic Saudou (U. Grenoble) for generously sharing huntingtin constructs, Amine Kamen (McGill U.) for guidance on CRISPR/Cas9, and the members of the Hendricks lab for thoughtful comments on the manuscript.

ENPP is supported by a CGS-M fellowship from the Natural Sciences and Engineering Council of Canada. AGH is supported by the Canadian Institutes for Health Research (PJT-159490).

## Notes

### Competing Interest Statement

The authors have declared no competing interest.

### Summary of Updates

The revised version contains updated analyses and experiments.

