## Supplemental Information for "Huntingtin S421 phosphorylation increases kinesin and dynein engagement on early endosomes and lysosomes"

### Supplementary materials

**Table S1. Calculated parameters from exponential fits for unbinding rates of early endosomes and lysosomes.**

| Condition | $U_0$ | $F_d$ (pN) |
|---|---|---|
| Early endosome WT outward | $2.1 \times 10^{-7}$ | 0.42 |
| Early endosome S421D outward | 0.31 | 3.2 |
| Early endosome WT inward | $7.2 \times 10^{-7}$ | 0.43 |
| Early endosome S421D inward | 0.31 | 6.3 |
| Lysosome WT outward | 0.49 | 12.1 |
| Lysosome S421D outward | 0.24 | 6.6 |
| Lysosome WT inward | 1.9 | 14.6 |
| Lysosome S421D inward | 0.77 | 10.8 |

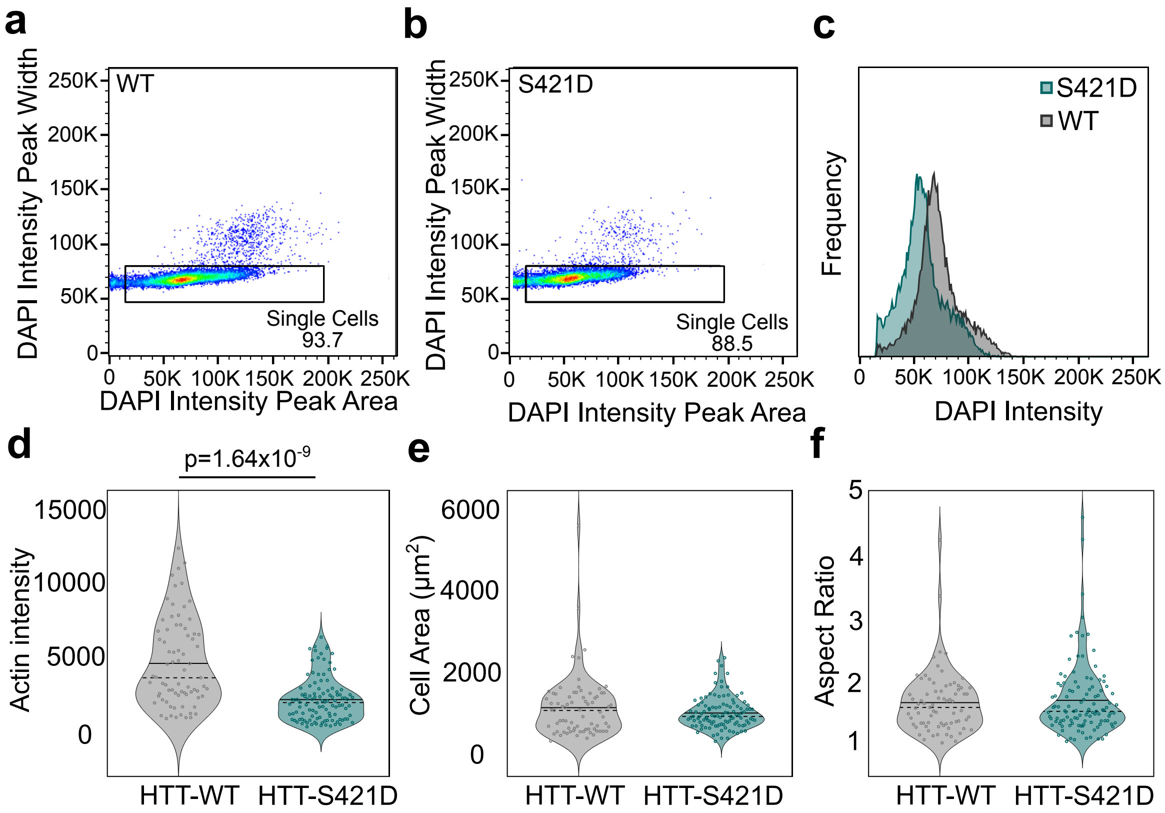


**Figure S1. HTT-S421D cell ploidy and morphology is not affected by the phosphomimetic mutation, however actin intensity is lower than HTT-WT.** (**a**,**b**) Fluorescence Associated Cell Sorting DAPI staining peak width versus area plot for HTT-WT cells (**a**) and HTT-S421D cells (**b**), with the heatmap indicating the density of datapoints with blue representing low density and red indicating high density. The box in the image indicates the threshold for single cells, with the number of single cells detected indicated on the image. (**c**) Distributions of the DAPI intensity for HTT-WT (grey), and HTT-S421D (teal) cells plotted on the same axes for comparison. (**d**) Distributions of actin intensities for HTT-WT and HTT-S421D cells as indicated. (**e**) Distributions of cell area in $\mu m^2$ for HTT-WT and HTT-S421D cells as indicated. (**f**) Distributions of cell aspect ratios for HTT-WT and HTT-S421D cells. Statistical significance by two-tailed Student t test is indicated by p values above the plot. Data from the same images as the actin immunofluorescence analysis in **Figure 1** with some additional images for the HTT-S421D condition. HTT-WT data was from 14 experiments, 84 cells, and HTT-S421D data from 11 experiments, 110 cells.

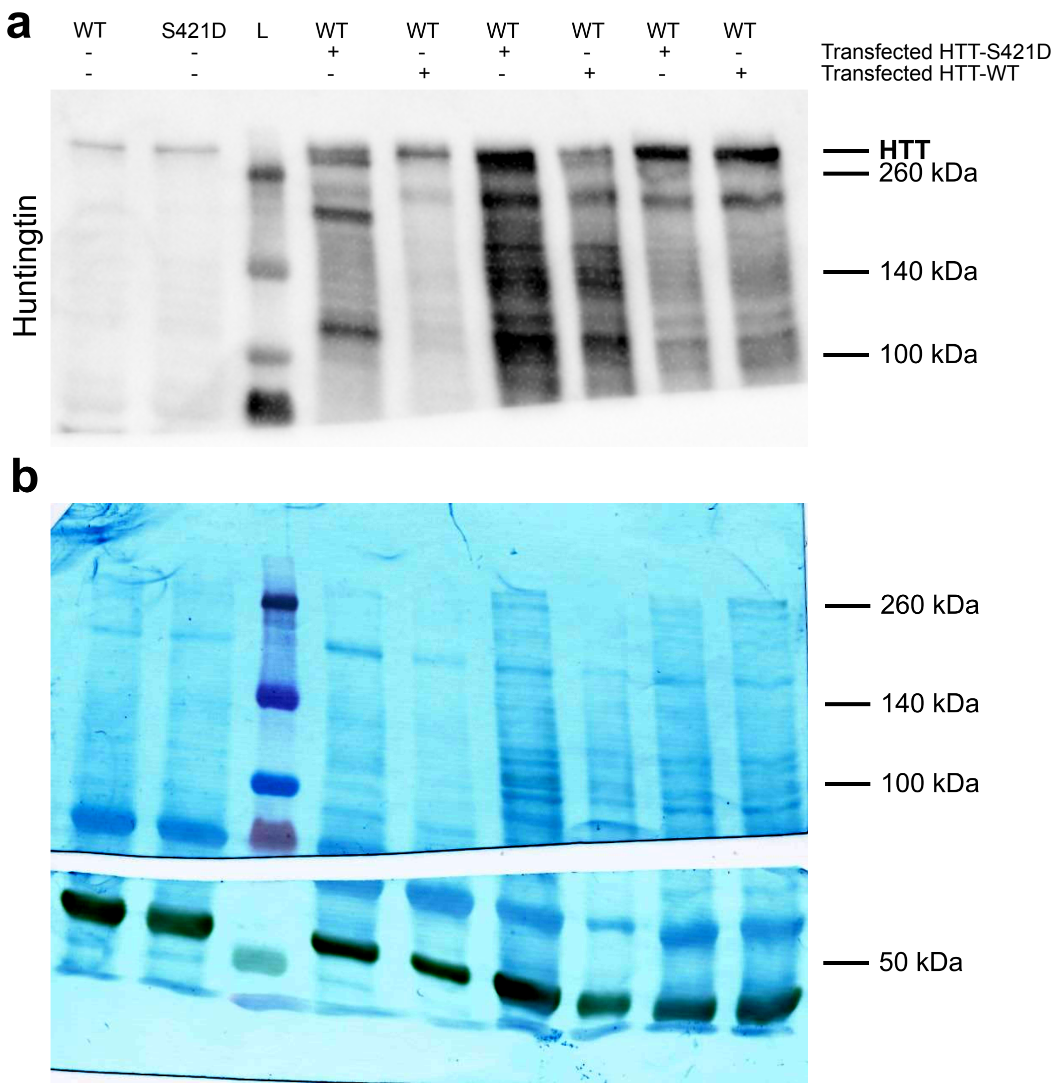


**Figure S2. Western blot transparency for huntingtin blot in Fig. 1.** (**a**) Full western blot (above 100 kDa) of the blot presented in Fig. 1G using the anti-huntingtin antibody. Ladder is indicated by L with molecular weights to the right of the blot. (**b**) Coomassie staining of the entire membrane post-antibody staining.

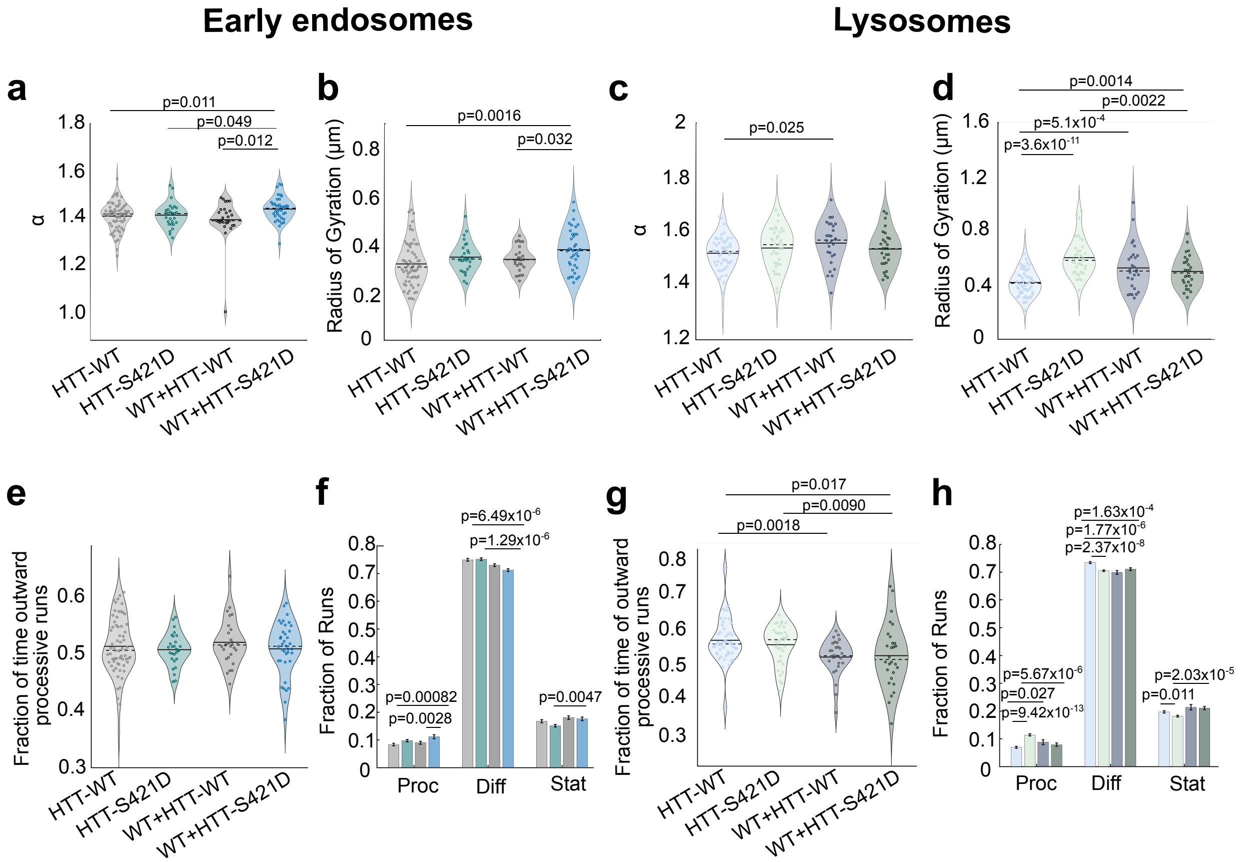


**Figure S3. Overexpression of HTT-WT or HTT-S421D affects all motility parameters for early endosomes and lysosomes.** (**a, b**) Early endosome (**a**) and lysosome (**c**) processivity per cell, measured by α, the slope of the log-log plot of their mean-squared displacement. Means are represented with a filled line, while medians are dashed lines. (**b,d**) Radius of gyration ($R_g$) per cell of early endosomes (**b**) and lysosomes (**d**). $R_g$ is a measure of the radius that contains half of the datapoints in the trajectory centered around its average position. (**e,g**) Early endosome (**e**) and lysosome (**g**) directional bias per cell of processive trajectories toward plus ends of microtubules (outwards), the remaining fraction of processive trajectories move inwards. Overexpression conditions represented by WT+HTT-WT or WT+HTT-S421D. Directional bias is determined by fraction of time particles are moving outwards compared to inwards and averaged for each cell. Note the endogenous expression data is the same as that in **Figure 2**. (**f, h**) Fraction of processive (Proc), diffusive (Diff), and stationary (Stat) runs for each condition in early endosomes (**f**) and lysosomes (**h**) with the same colour code as previous images. For early endosomes (**f**), HTT-WT is light grey, HTT-S421D is teal, WT+HTT-WT is dark grey, and WT+HTT-S421D is blue. For lysosomes (**h**), HTT-WT is light blue, HTT-S421D is light green, WT+HTT-WT is navy and WT+HTT-S421D is dark green. The number of experiments, cells, and trajectories for each condition are the following; HTT-WT early endosomes: 65 cells, 41994 trajectories, 6 experiments; HTT-S421D early endosomes: 28 cells, 25476 trajectories, 3 experiments; WT+HTT-WT early endosomes: 28 cells, 17424 trajectories, 5 experiments, WT+HTT-S421D early endosomes: 39 cells, 22854 trajectories, 6 experiments; HTT-WT lysosomes: 55 cells, 25247 trajectories, 5 experiments; HTT-S421D lysosomes: 38 cells, 12670 trajectories, 4 experiments; WT+HTT-WT lysosomes: 29 cells, 7558 trajectories, 5 experiments; WT+HTT-S421D lysosomes: 28 cells, 7031 trajectories, 6 experiments. Statistical significance was determined via Student's two-tailed t-test and p-values are indicated on each plot where applicable.

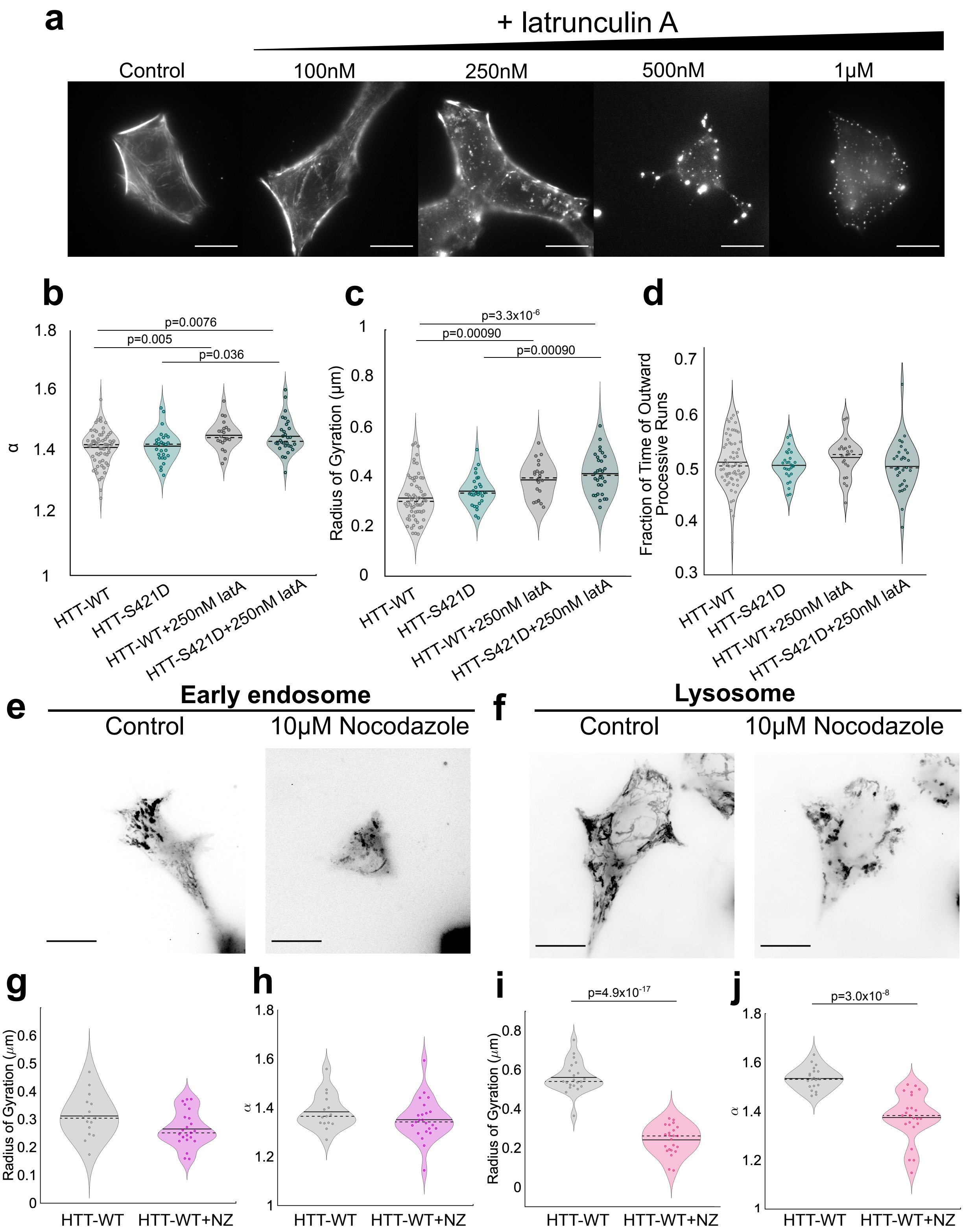


**Figure S4. Cytoskeletal disruption affects cargo motility.** (**a**) Actin cytoskeleton labelled with SiR actin in HTT-S421D HEK293T cells treated with 100 nM-1 μM concentrations of latrunculin A (latA). Scale bars for all images are 20 μm, all images inverted. (**b**) Per cell α values of Rab5-eGFP labelled early endosomes. Means are represented with a filled line, while medians are dashed lines. (**c**) Radius of gyration (Rg) per cell is presented for early endosomes. (**d**) Directional bias per cell shown by fraction of time processive runs moving outward for early endosomes. Note: control HTT-WT and HTT-S421D data for latrunculin A experiments is the same data reported in Figure 2. (**e, f**) Maximum projections of a cell before and after nocodazole (NZ) treatment for early endosome and lysosome analysis. (**g-j**), Radius of gyration and alpha values for early endosomes and lysosomes before after nocodazole treatment. (**b-d**) HTT-WT control data from 7 independent experiments (41994 trajectories, 64 cells), HTT-S421D data from 3 independent experiments (25476 trajectories, 28 cells), HTT-WT+latA data from 3 independent experiments (17461 trajectories, 22 cells); HTT-S421D+latA data from 5 independent experiments (19879 trajectories, 30 cells). (**g, h**) HTT-WT control data from 4 experiments, 22 cells, 6109 trajectories and WT+nocodazole data is from 4 experiments, 23 cells, and 7094 trajectories. (**i, j**) Contains control data from 4 experiments, 15 cells, 17073 trajectories, and nocodazole treated data from 4 experiments, 23 cells, and 6639 trajectories. Statistical significance of $p<0.05$ by Student's two sample t-test indicated in the figure where applicable.

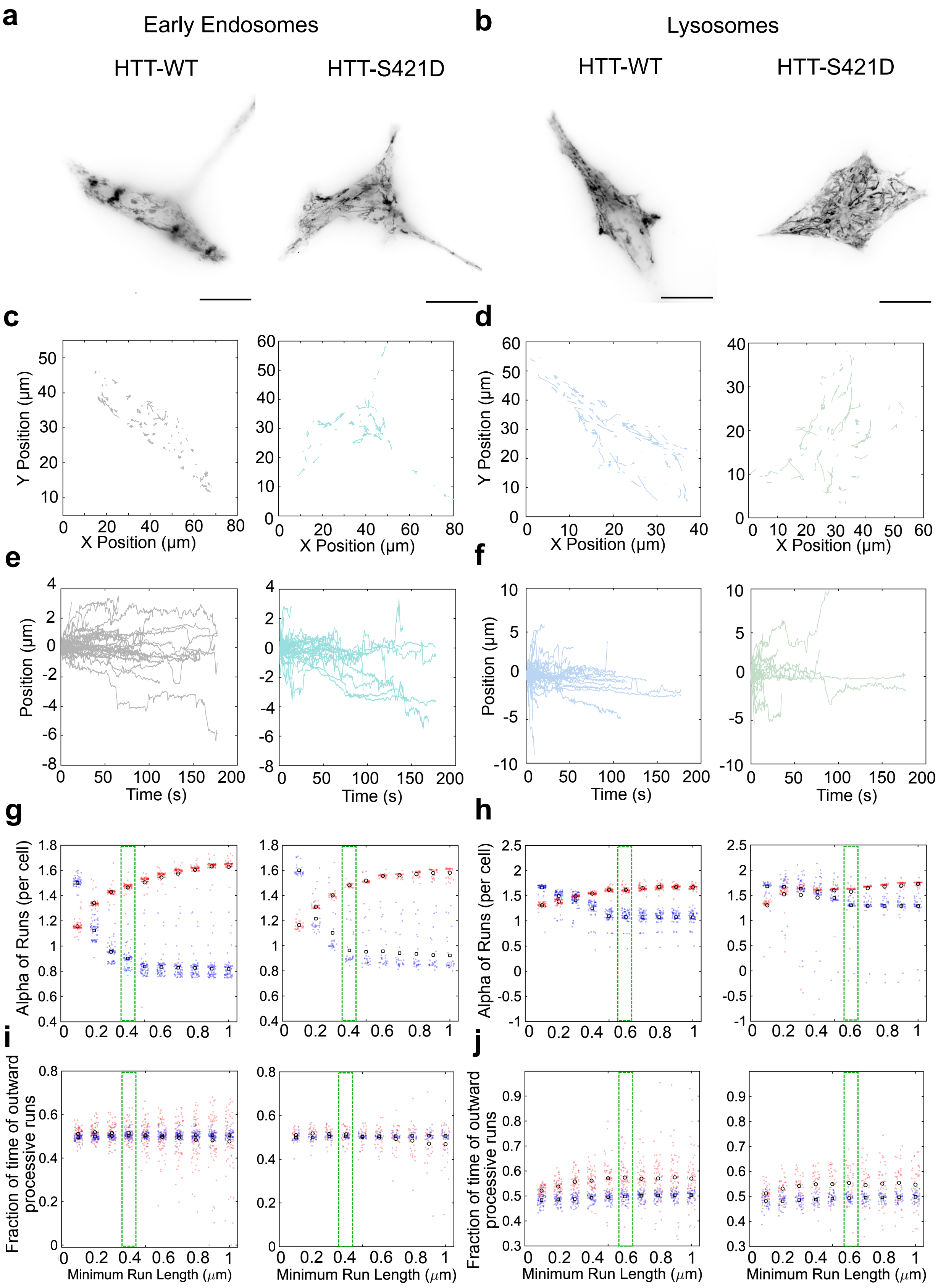

a
Early Endosomes
HTT-WT
HTT-S421D
b
Lysosomes
HTT-WT
HTT-S421D
c
d
Y Position (μm)
X Position (μm)
e
f
Position (μm)
Time (s)
g
h
Alpha of Runs (per cell)
i
j
Fraction of time of outward processive runs
Minimum Run Length (μm)

**Figure S5. Representations of single cell trajectories and minimum run length for processive runs analysis.** (**a, b**) Additional maximum projections of early endosomes (**a**) and lysosomes (**b**) for HTT-WT and HTT-S421D. (**c,d**) Approximately 100 trajectories from the cells directly above in (**a,b**) plotted in the xy positions as detected using TrackMate. (**e,f**) Position vectors generated using the dot product of the trajectories from (**c,d**) and centered at 0 with inward trajectories having negative values and positive values indicating outward trajectories. (**g,h**) Alpha of processive (red) and diffusive (blue) runs for early endosomes (**g**) and lysosomes (**h**) depending on the minimum run length for processive runs on the x axis. The green dashed box indicates the minimum run length of processive runs chosen for analysis of early endosomes and lysosomes respectively based on these plots. (**i,j**) Fraction of outward processive (red) and diffusive (blue) runs for early endosomes (**i**) and lysosomes (**j**) with the minimum run length of processive runs as indicated on the x axis. The green box indicates the chosen minimum run length of processive runs based on the alpha plots above for all trajectory analyses.

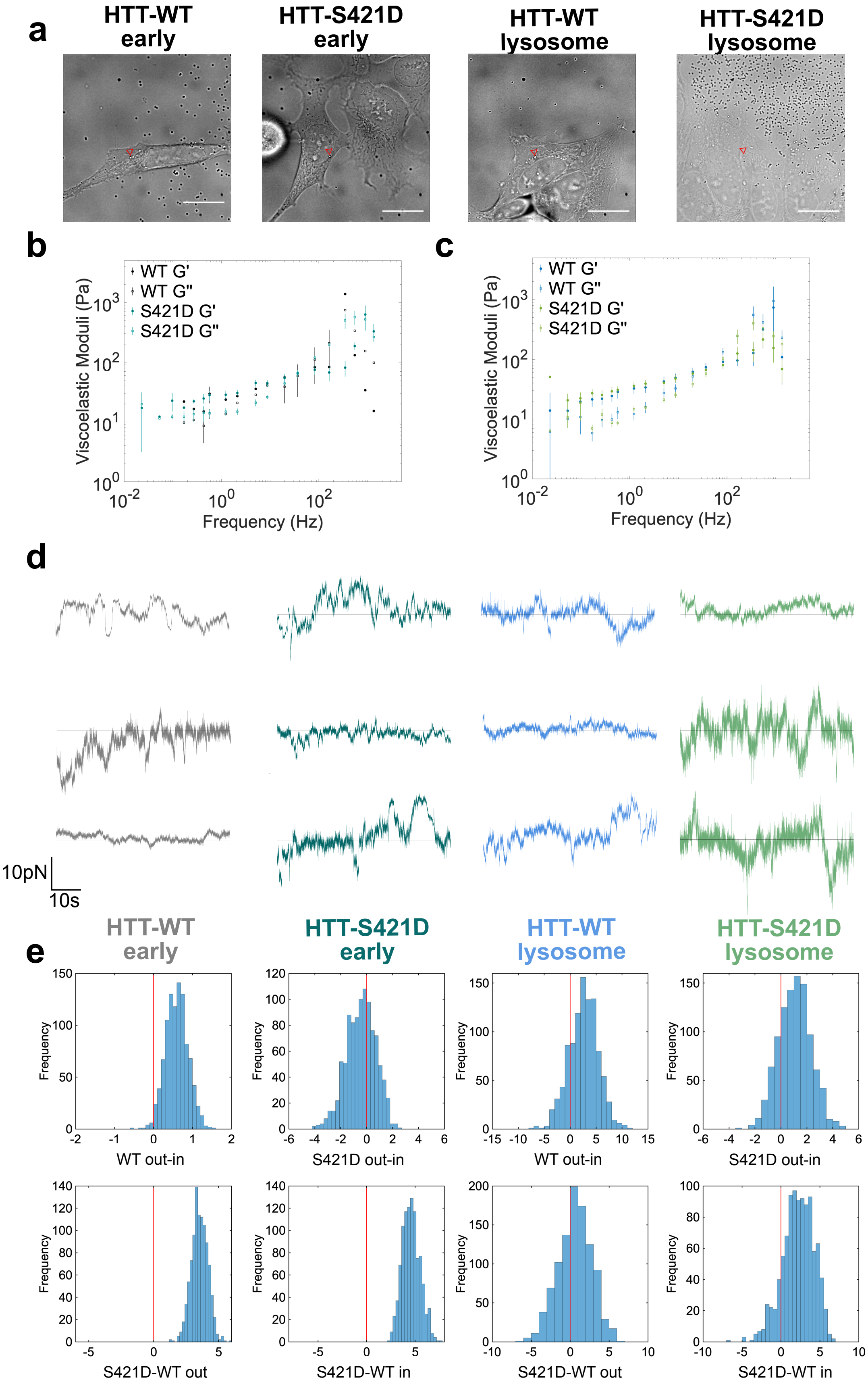

a
HTT-WT early
HTT-S421D early
HTT-WT lysosome
HTT-S421D lysosome
b
WT G'
WT G''
S421D G'
S421D G''
Viscoelastic Moduli (Pa)
Frequency (Hz)
c
WT G'
WT G''
S421D G'
S421D G''
Viscoelastic Moduli (Pa)
Frequency (Hz)
d
10pN
10s
HTT-WT early
HTT-S421D early
HTT-WT lysosome
HTT-S421D lysosome
e
Frequency
WT out-in
S421D out-in
WT out-in
S421D out-in
S421D-WT out
S421D-WT in
S421D-WT out
S421D-WT in

**Figure S6. Force measurement analysis approach, additional force traces, and viscoelastic properties of HTT-WT and HTT-S421D cells.** (**a**) Brightfield image of HTT-WT and HTT-S421D cells containing a trapped phagocytosed 500 nm bead (red arrow) during measurement of early endosome (early) or lysosome forces. (**b, c**) Viscous (G'') and elastic moduli (G') for HTT-WT (black) and HTT-S421D (teal) early endosomes and HTT-WT (blue) and HTT-S421D (green) lysosomes. (**d**) Additional sample force traces for each condition in optical tweezers assays, the horizontal line indicates the zero axis. Positive value forces are directed outward and negative forces are directed inward. (**e**) Distributions of bootstrap mean differences evaluated for additional statistical analysis of force distributions for the differences shown on the x axis of each plot in both early (left) and late (right) endosomes.

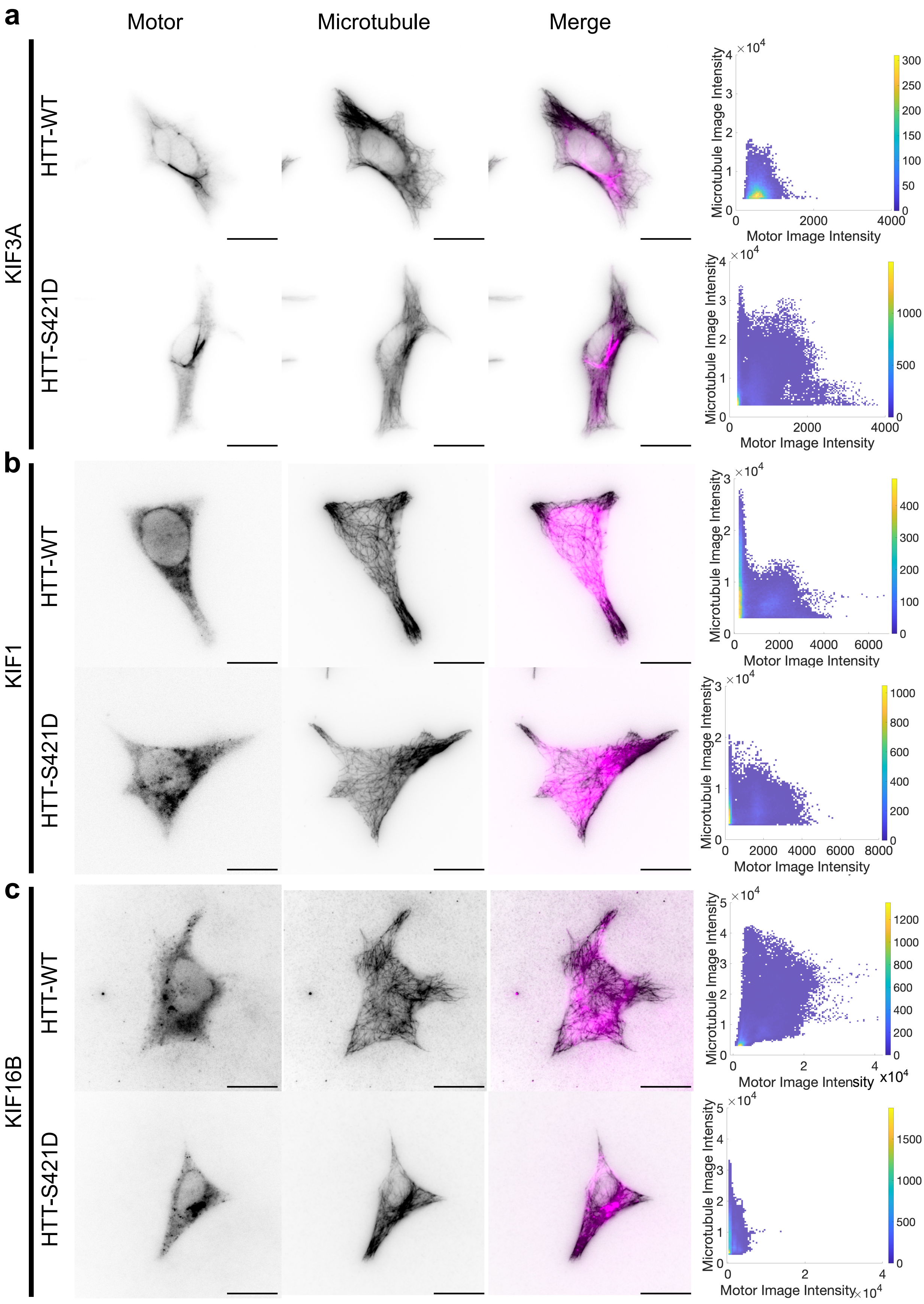
a
b
c
Motor
Microtubule
Merge
KIF3A
KIF1
KIF16B
HTT-WT
HTT-S421D
Microtubule Image Intensity
Motor Image Intensity
×10^4
x10^4

**Figure S7. HTT-S421D recruits less kinesin-2 and -3 to microtubules.** (**a**), Inverted immunofluorescence images demonstrating colocalization (rightmost image) with kinesin in magenta and microtubules in black, kinesin-2 (kif3a, left), on extracted microtubules (middle) for both HTT-WT (top) and HTT-S421D (bottom) cells. Colocalization plots were created from plotting intensity values from both motor and microtubule channels of all images (right). All scale bars are 20 μm. (**b**) The kif1 form of kinesin-3, with the data organized the same as in (**a**). (**c**) The kif16b form of kinesin-3, with the data organized as in (**a**). For kif1, there were no significant differences in colocalization. For kif3a and kif16b, cells expressing HTT-WT had higher colocalization with p values of $1.85 \times 10^{-6}$ and $1.77 \times 10^{-5}$, respectively by Student's t-test. HTT-WT kif3a data from 29 cells, 3 experiments, HTT-S421D kif3a data from 32 cells, 3 experiments, HTT-WT kif1adata from 13 cells, 2 experiments, HTT-S421D kif1 data from 19 cells, 2 experiments, HTT-WT kif16b data from 15 cells, 2 experiments, and HTT-S421D kif16b data from 18 cells, 2 experiments.
